# Heat Shock Transcription Factor B1 Coordinates Distinct bHLH Modules to Orchestrate Land Plant Adaptation

**DOI:** 10.64898/2026.11.09.765476

**Authors:** Huan-Chi Tien, Kuan-Hung Lin, Lavakau Thalimaraw, Ming-Ren Yen, Hsiu-Ju Yang, Ya-Ru Li, Chia-Yi Cheng, Ming-Jung Liu, Ting-Ying Wu

**Author notes:** These authors contribute equally to this study (Huan-Chi Tien, Kuan-Hung Lin). Correspondence: Ting-Ying Wu, Institute of Plant and Microbial Biology Academia Sinica, Taipei, Taiwan.

## Abstract

Adaptation to terrestrial environments required the evolution of morphological structures, such as rhizoids and air pores, and robust tolerance mechanisms to environmental stress. While these innovations are well-described, the regulatory logic coordinating development with stress signaling during this transition remains largely unknown. Here, we identify Heat Shock Transcription Factor B1 (HSFB1) as a central evolutionary node that bridges this gap in *Marchantia polymorpha*. Using single-nucleus RNA sequencing (snRNA-seq) and gene regulatory network analysis, we reveal that MpHSFB1 is required to activate cell-type-specific programs for terrestrial adaptation, functioning via the regulation of cytokinin and reactive oxygen species (ROS) homeostasis. We find that MpHSFB1 functions through a modular logic: it co-opts a specific set of bHLH transcription factors (including MpbHLH21/40) to drive developmental morphogenesis, while recruiting a distinct partner (MpbHLH51) to confer resistance to prolonged heat stress. This functional uncoupling allows a single ancestral regulator to manage disparate physiological needs. By reconstructing HSFB1 regulons across diverse plant lineages, we propose that MpHSFB1 acted as an ancestral pleiotropic hub that was effectively rewired and subfunctionalized in later angiosperm lineages for land plant evolution.

## Introduction

Land plants underwent several key morphological modifications to adapt to terrestrial environments. One such innovation was the development of rhizoids—root-hair-like structures that arise from epidermal cells to facilitate water and nutrient absorption. Another was the formation of air pore complexes on the plant surface, which enable gas exchange while minimizing water loss^1^. In addition, land plants evolved mechanisms to cope with oxidative stress caused by factors such as heat, water, nutrient scarcity, and UV radiation. These mechanisms involve the perception of stressors and activation of signal transduction pathways that include reactive oxygen species (ROS), specific phytohormones, and the co-option of multiple transcription factors (TFs)^2^. The integration of these processes requires the coordination of both internal and external signals^3^. Consequently, understanding how plants perceive and process environmental cues through multilayered molecular networks, and how these networks influence development and survival strategies, remains a central focus of research on land plant adaptation.

One such group of phytohormones, cytokinin, plays a crucial role not only in plant development but also in responses to abiotic and biotic stresses. Bryophytes already possess a complete cytokinin signaling system, in which the signal is perceived and transmitted through a variant of the Two-Component System (TCS), which consists of CHASE domain–containing histidine kinase receptors (CHKs), histidine-containing phosphotransfer proteins (HPTs), and response regulators (RRs)^4,5^. Although this primitive signaling pathway originated in green algae, it acquired pleiotropic functions during land colonization by integrating with other signaling and gene regulatory modules, enabling the coordination of growth, development, and stress responses under terrestrial conditions^6,7^. Consequently, mutants defective in cytokinin signaling exhibit diverse morpho-physiological phenotypes across land plants, ranging from bryophytes to angiosperms, including altered root and gametophyte development and increased sensitivity to drought or salt stress^8^. However, the specific TFs that integrate these developmental and environmental cues to facilitate adaptation remain largely unresolved.

While algal lineages possess ancestral Heat Shock Transcription Factor (HSF), the distinct divergence into class A and class B HSFs first occurred in the ancestral land plant*^9,10^*. Unlike HSFA1, which serves as the conserved master regulator of high-temperature responses in plants, HSFB1 acquired more diversified functions^11-13^. For example, Mp*hsfb1* knockout mutants in *Marchantia* exhibit only minor sensitivity to high temperatures*^9^*. In contrast, the development of air pores, air chambers, and rhizoids was defective in *Marchantia* Mp*hsfb1* mutants, resembling the defects seen in mutants with impaired cytokinin signaling^14-16^. Given the involvement of HSFB1 in traits related to morphological adaptation to the terrestrial environment, we hypothesize that MpHSFB1 (Mp4g12230) evolved as a central regulator that balances external cues (e.g., temperature) with internal hormonal signaling pathways (e.g., cytokinin). This coordination likely orchestrates developmental responses and defense strategies necessary to overcome the challenges of life on land. However, the underlying mechanisms by which HSFB1 achieves such precise spatiotemporal regulation under various conditions remain unclear.

Basic helix–loop–helix (bHLH) proteins are ancient TFs found across all eukaryotes and represent one of the most expansive and evolutionarily versatile TF families in plants^17^. Plant bHLHs form a monophyletic group distinct from those in rhodophytes (red algae) and metazoans^18^. A major radiation of the bHLH family occurred after the divergence of the moss and vascular plant lineages, giving rise to approximately 26–32 subfamilies that largely emerged during the critical phase of land colonization^19^. This expansion likely provided the regulatory versatility needed for plants to evolve novel organs and developmental plasticity in fluctuating terrestrial environments. bHLHs commonly form homo- or heterodimers and function as transcriptional activators and/or repressors of genes involved in key developmental processes and their integration with environmental cues such as high temperature and nutrient deficiency^17^. For example, the bHLHs ICE1/2, SPCH, MUTE, and FAMA form heterodimers to regulate stomatal development, while LRL1/2 and RSLs are involved in root hair formation in *Arabidopsis*^20-22^. This developmental role extends to bryophytes, where ICE1 (MpbHLH16, Mp4g04910) and ZOU1 (MpbHLH38, Mp4g19650) promote pegged rhizoid differentiation, while RSL1 (MpbHLH14, Mp3g17930) drives general rhizoid formation^23,24^. In addition, illustrating their role in environmental integration, the bHLH PIF4 regulates SPCH in response to high ambient temperatures in *Arabidopsis*^25^. Yet, how these bHLHs act synergistically or antagonistically to regulate specific downstream targets, despite sharing similar DNA-binding sites, remains an open question. One plausible explanation is that they cooperate with distinct TFs in specific regulatory contexts. Elucidating the combinatorial regulation between bHLHs and other TFs will provide crucial insights into how the diversification of transcriptional networks contributed to developmental complexity during land plant evolution.

The ability to integrate these pathways is key to survival and colonization on land. In this study, we demonstrate that HSFB1 functions as an ancestral regulatory hub that maintains homeostasis between development and stress responses through modular interactions with different bHLH TFs. Through integrated analyses of single-nucleus RNA-seq, bulk time-course RNA-seq, and phenotypic observations using various genetic mutants, we found that MpHSFB1 and MpbHLHs regulate early developmental processes in *Marchantia* via cytokinin. In contrast, MpHSFB1 modulates a distinct set of MpbHLHs to confer tolerance during prolonged heat stress. Furthermore, swapDAP-seq analysis revealed that HSFB1 spatiotemporally regulates diverse gene functions across land plants. Together, these findings highlight how HSFB-mediated diversification contributed to the evolution of complex developmental and stress-response networks in plants.

## Results

### MpHSFB1 regulates morphological adaptations associated with terrestrial lifestyles in *Marchantia*

*Marchantia* evolved various morphological and physiological adaptations to overcome unfavorable environmental conditions. These structures included the development of air pores, rhizoids, and mechanisms for hormone homeostasis and ROS regulation (Figure 1A). We observed developmental defects in the Mp*hsfb1* knockout mutant during the early developmental stage of *Marchantia*, starting from day 7 (Figure 1B, Supporting Figure S1, Supporting Dataset S1). At day 14, the Mp*hsfb1* mutant exhibited several distinct morphological abnormalities compared to wild-type Tak-1 plants. Consistent with previously reported phenotypes^9,26^, these included a ∼50% higher rhizoid-to-thallus density, an absence of air pore and air chamber formation, and an abnormally upward-curved thallus. In addition to these known traits, we revealed several novel phenotypic characteristics in the Mp*hsfb1* mutant, including a ∼3.5-fold smaller surface area, a ∼1.3-fold lower solidity value, and ∼1.8-fold higher ROS levels (Figures 1C and 1D, Supporting Figure S1, Supporting Dataset S1). In contrast, the Mp*HSFB1*ox transgenic lines showed no differences in air pore and air chamber formation, thallus area, or rhizoid-to-thallus ratio when compared to Tak-1 plants. However, they exhibited significantly higher solidity (∼1.2-fold) and lower ROS levels (∼1.3-fold) (Figures 1C and 1D, Supporting Figure S1, Supporting Dataset S1). Together with the evolution of HSFB1 in the ancestral land plant lineage^9,10^ (Supporting Figure S1C), we hypothesize that MpHSFB1 plays a crucial role in regulating terrestrial adaptation. To test this, we designed a comprehensive study integrating time-course RNA-seq, snRNA-seq, and DAP-seq analyses in Tak-1 (wild-type) plants, as well as in Mp*hsfb1* and Mp*HSFB1ox* transgenic lines. These molecular data were combined with time-series phenotypic observations to capture the regulatory roles of MpHSFB1 across different developmental stages and cell types in *Marchantia* (Figure 1E).

**Figure 1.**
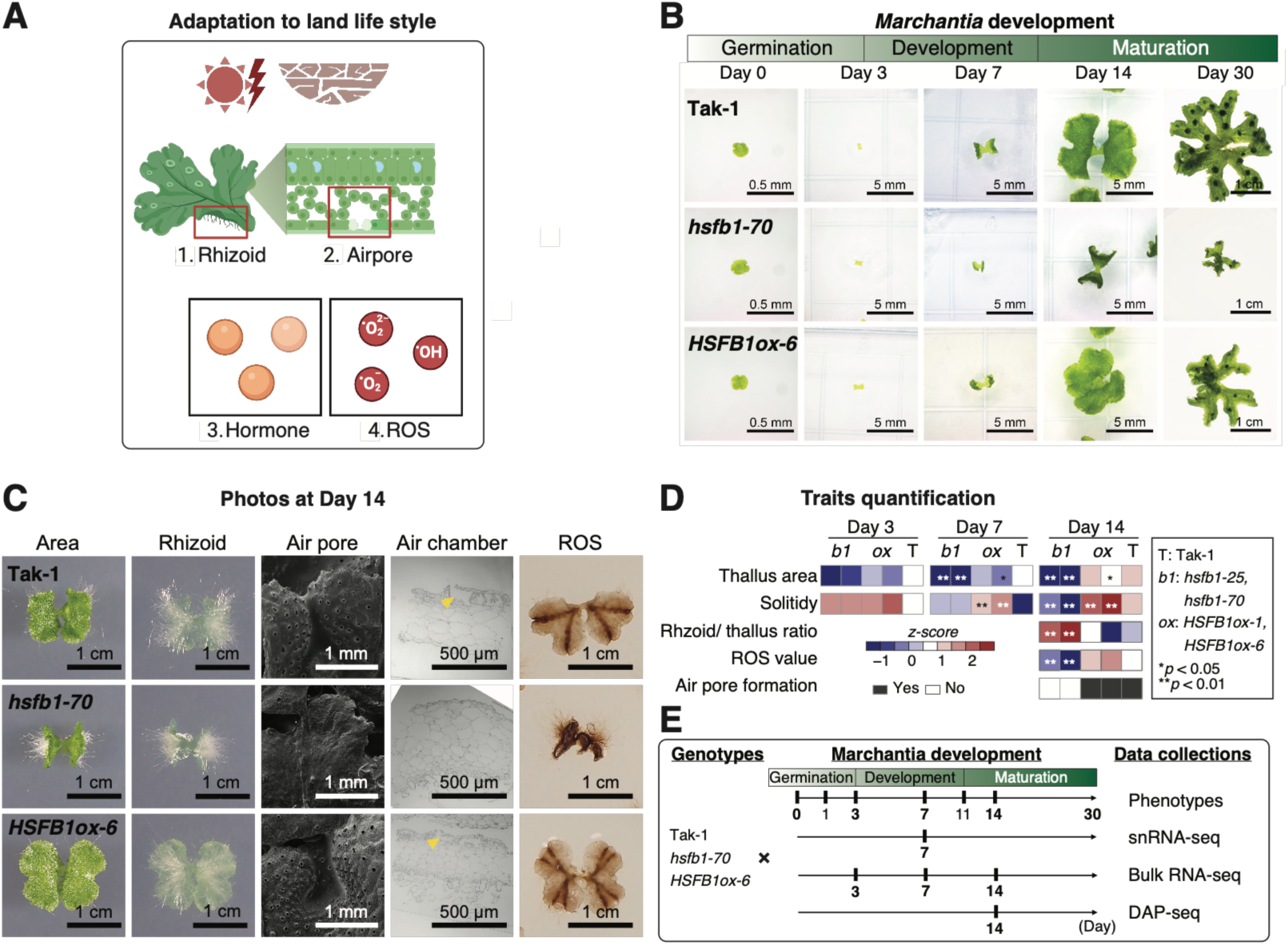
MpHSFB1 is involved in terrestrial adaptation. **A.** Schematic illustration of four key features associated with adaptation to terrestrial life. 1. Plants evolved rhizoid structures for water and nutrient absorption. 2. They also developed air pore structures for gas exchange. 3–4. In addition, hormonal and ROS signaling pathways evolved to enable responses to unfavorable environments. B. Representative images of Tak-1, Mp*hsfb1*, and Mp*HSFB1ox* plants from day 0 to day 30. **C.** Representative images showing developmental traits at day 14. **D.** Heatmaps showing quantitative analyses of developmental traits. Z-scores were calculated across developmental stages and genotypes. p-values were determined using Student’s t-test between mutants and the wild type. Two independent experiments were performed with *n* = 30–60 per time point. The raw measurements and complete statistical results are provided in Supporting Dataset S1. **E.** Schematic overview of the datasets collected in this study.

### MpHSFB1 influences cell type abundance and similarity in *Marchantia*

To investigate cellular-level changes in Mp*hsfb1* mutants, we generated single-nucleus RNA-seq (snRNA-seq) datasets from Tak-1 plants and Mp*hsfb1* mutants using the 10x Genomics platform (Supporting Figure S2A, Supporting Dataset S2). In total, we collected over 20,000 nuclei per genotype. After quality control filtering, we retained 18,073 and 14,853 high-quality nuclei from Tak-1 and Mp*hsfb1*, respectively. The datasets showed comparable depth, with a median of ∼2,060 unique transcripts and ∼1,150 detected genes per nucleus for both genotypes (Figure 2A, Supporting Dataset S2).

**Figure 2.**
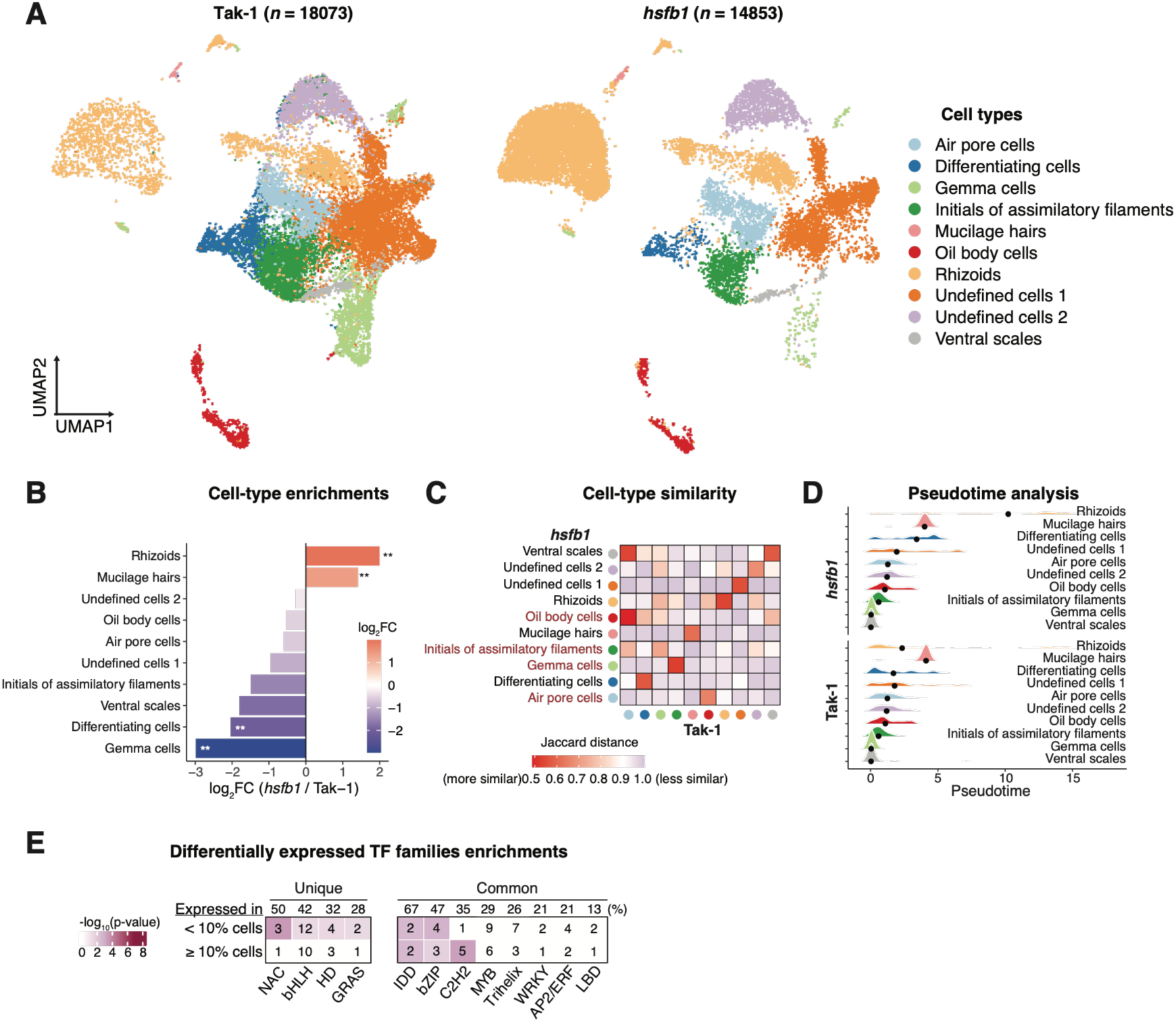
snRNA-seq reveals cell-type changes in Mp*hsfb1* mutants. **A.** UMAP visualization of major cell-type clusters in Tak-1 plants and Mp*hsfb1* mutants. **B.** Bar graph showing cell-type enrichments between Tak-1 and Mp*hsfb1*. **p < 0.01, hypergeometric test. **C.** Heatmap showing cell-type similarity between Tak-1 and Mp*hsfb1*. The color key represents Jaccard distance, where lower values indicate higher similarity. **D.** Ridgeline plot showing pseudotime distributions across cell types in Tak-1 and Mp*hsfb1*. **p < 0.01, empirical cumulative distribution function (ECDF), Kolmogorov–Smirnov test. **E.** Heatmap showing transcription factor (TF) families differentially expressed (DE) between Tak-1 and Mp*hsfb1*, enriched in high (>10%) or low (<10%) proportions of cells. Percentages were calculated as the number of DE TFs over the total TFs within each family. Note that only the TF families with a number of DE TFs greater than 2 were shown. The full list was presented in the Supporting Dataset S2.

We clustered these nuclei into distinct cell types based on a previously published *Marchantia* reference cell atlas^26^ (Supporting Figure S2B, Supporting Dataset S2). Most major cell types were successfully resolved and comparable to the protoplast-based scRNA-seq atlas. We observed significant shifts in cell type abundance: rhizoid and mucilage hair cells were significantly increased in the mutant, whereas gemma and differentiating cells were markedly reduced (Figure 2B, Supporting Figure S3, Supporting Dataset S2). This cellular composition shift mirrors the observed macroscopic phenotypes.

Furthermore, cell-type marker expression analysis revealed that air pore- and oil body-specific genes exhibited the greatest transcriptional divergence between Tak-1 and Mp*hsfb1*, suggesting that cell-type identity is compromised in the mutant (Figure 2C, Supporting Dataset S2). Consistent with this transcriptomic profile, histochemical staining confirmed that Mp*hsfb1* mutants possess a significantly reduced number of oil bodies (Supporting Figure S3D & S3E). Pseudotime analysis indicated that the developmental trajectories of rhizoid and differentiating cells were significantly altered in Mp*hsfb1* compared to Tak-1 (Figure 2D, Supporting Dataset S3). Finally, we examined the differentially expressed transcription factor (TF) families between Tak-1 plants and Mp*hsfb1* mutants. We found that bHLH (*n* = 12, p < 0.01) and NAC (*n* = 3, p < 10^-4^) TFs were among the most highly ranked families in both unique and common cell populations (Figure 2E, Supporting Figures S3B & S4, Supporting Dataset S3). These results suggest that MpHSFB1 regulates development in *Marchantia* in a cell-type-specific manner, likely through the co-regulation of bHLH and NAC transcription factors.

### The Mp*hsfb1* regulons underwent functional rewiring through altered gene targeting

To further explore regulation at the cellular level, we constructed gene regulatory networks (GRNs) to dissect the distinct and overlapping regulons in each genotype. Although ∼75% of the regulons (*n* = 32) were shared between the two genotypes, the networks from wild-type Tak-1 plants displayed a denser, more interconnected structure. Specifically, Tak-1 networks contained ∼1.3-fold more nodes and exhibited ∼1.5-fold higher average connectivity compared to those in the Mp*hsfb1* mutants (Figure 3A, Supporting Figures 5A-5C, Supporting Dataset S4). This suggests that the Mp*hsfb1* mutant has a fragmented regulatory system, leading to uncoordinated development and disrupted stress homeostasis.

**Figure 3.**
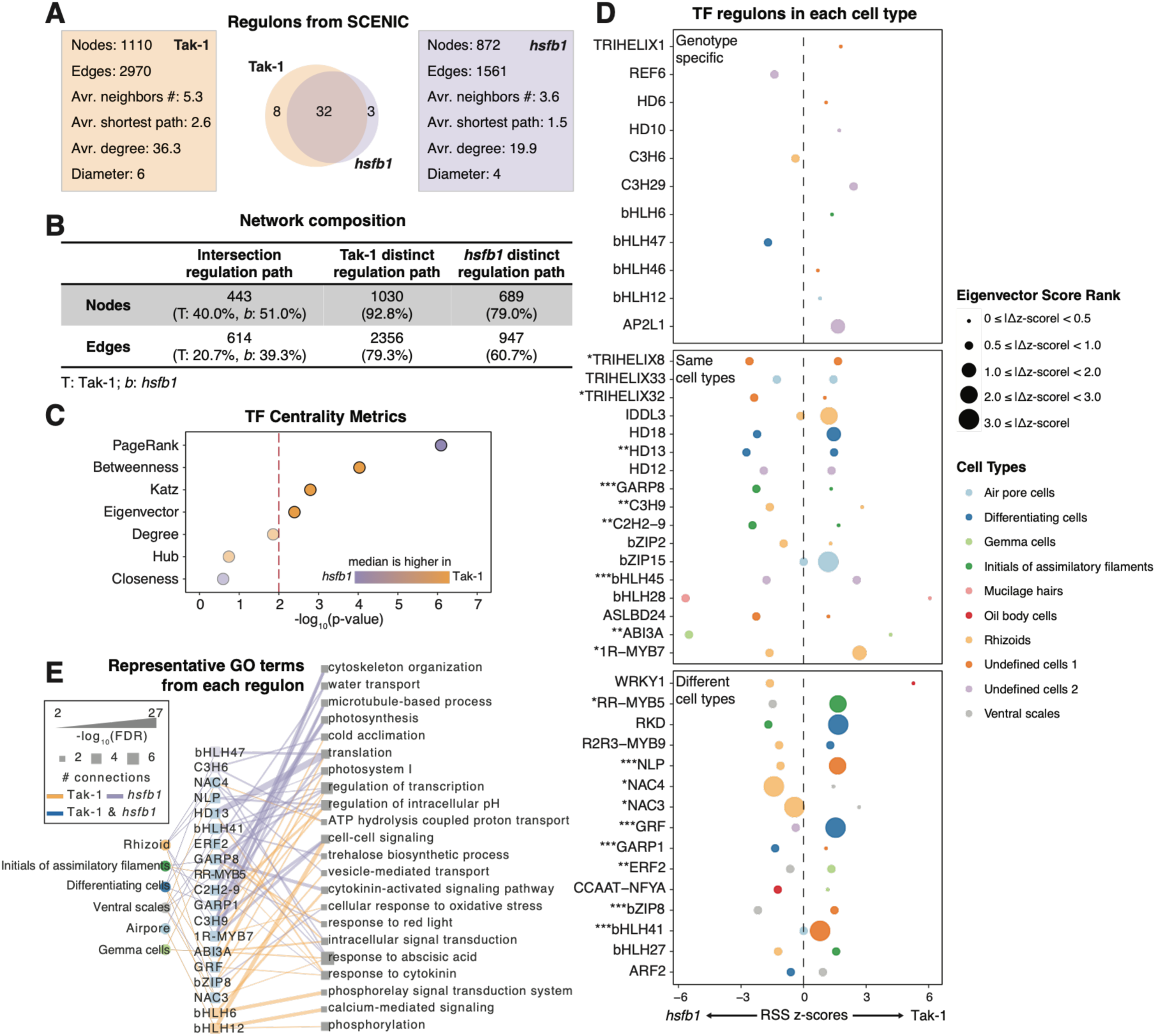
The Mp*hsfb1* regulons underwent functional rewiring through altered gene targeting. **A.** Comparison of regulon properties between Tak-1 and Mp*hsfb1*. The Venn diagram shows the number of overlapping regulons between the two genotypes. **B.** Table comparing the network composition of Tak-1 and Mp*hsfb1*. **C.** Dot plot showing the enrichment (hypergeometric test) of TF centrality metrics between Tak-1 and Mp*hsfb1*. **D.** Dot plots represent the scaled regulon specificity score (RSS) after z-score transformation in Tak-1 and Mp*hsfb1*. Dot size indicates the Δz-score between Tak-1 and Mp*hsfb1*, and dot color denotes the cell type with the maximum RSS for each regulon. **E.** Network visualization showing representative GO terms from each regulon in selected cell types. Edge width represents the significance of GO term enrichment, and node size indicates the number of connected regulons (TFs).

This structural difference was further supported by the observation that, while common nodes accounted for 40% and 51% of the total nodes in Tak-1 and Mp*hsfb1* regulons, respectively, the Tak-1-specific regulons contained a significantly greater number of nodes and edges (Figure 3B, Supporting Figure S5C, Supporting Dataset S4). Additionally, the TF nodes in Tak-1 regulons displayed significantly higher betweenness, Katz, and eigenvector centralities. This suggests that in the wild type, these TFs are highly connected and play more integrative roles across the network compared to the mutant (Figure 3C, Supporting Dataset S4).

We next examined the contribution of each regulon across cell types by ranking them using the regulon specificity score (RSS). Among the 32 common regulons, 17 (53.1%) retained their cell-type specificity, exhibiting the highest RSS rank in the same cell type in both genotypes. However, 15 (46.9%) showed shifted specificity, ranking highest in different cell types between the two genotypes (Figure 3D, Supporting Figure S5B, Supporting Dataset S4).

Gene Ontology (GO) analysis further revealed that the biological functions of these regulons diverged significantly. The regulons in Mp*hsfb1* mutants were primarily associated with housekeeping and structural functions such as photosynthesis, translation, and cell wall organization. In contrast, regulons in Tak-1 plants were enriched for functions linked to environmental adaptation, including cytokinin signaling and oxidative stress responses (Figure 3E, Supporting Figure S3C, Supporting Dataset S5). These results suggest that while the core regulatory architecture is partially conserved, a substantial fraction of regulons has been functionally reassigned in the absence of MpHSFB1, indicating HSFB1-dependent rewiring of cell-type-specific regulatory programs.

### MpHSFB1 and MpbHLHs regulate cytokinin homeostasis during the early *Marchantia* development

Developmental bulk RNA-seq at days 3, 7, and 14 identified 4,011 differentially expressed genes (DEGs) across eight clusters (C1-C8) (Figure 4A, Supporting Dataset S6). In Mp*HSFB1ox* transgenic lines, cluster C7 genes peaked at day 7 and were significantly enriched for cytokinin response functions (q-value < 10⁻⁸) (Figure 4B, Supporting Figures S6A & S6B). Consistent with this, cytokinin biosynthesis genes were differentially expressed between Tak-1 and Mp*hsfb1* mutants across the developmental time course and within specific tissues, including air pore cells, gemma cells, and ventral scales from the snRNA-seq analysis (Supporting Figures S2-S4 & S6C, Supporting Dataset S2 & S6). To determine whether this transcriptional signature was associated with altered cytokinin homeostasis, we measured endogenous active cytokinin (*cis*-zeatin) levels. *Cis*-zeatin concentrations were significantly lower (∼2-fold, *p* = 0.001) in Mp*hsfb1* mutants compared to Tak-1, but approximately 2- to 7-fold higher in Mp*HSFB1ox* transgenic lines across all time points (Figure 4C, *p* = 0.007). Furthermore, exogenous cytokinin application rescued the Mp*hsfb1* developmental phenotypes, restoring both thallus area (∼2-fold increase) and thallus angle (∼1.3-fold increase) to near-wild-type levels (Figure 4D & 4E, Supporting Dataset S1). Together, these results indicate that MpHSFB1 regulates early *Marchantia* development, at least in part, through cytokinin homeostasis.

**Figure 4.**
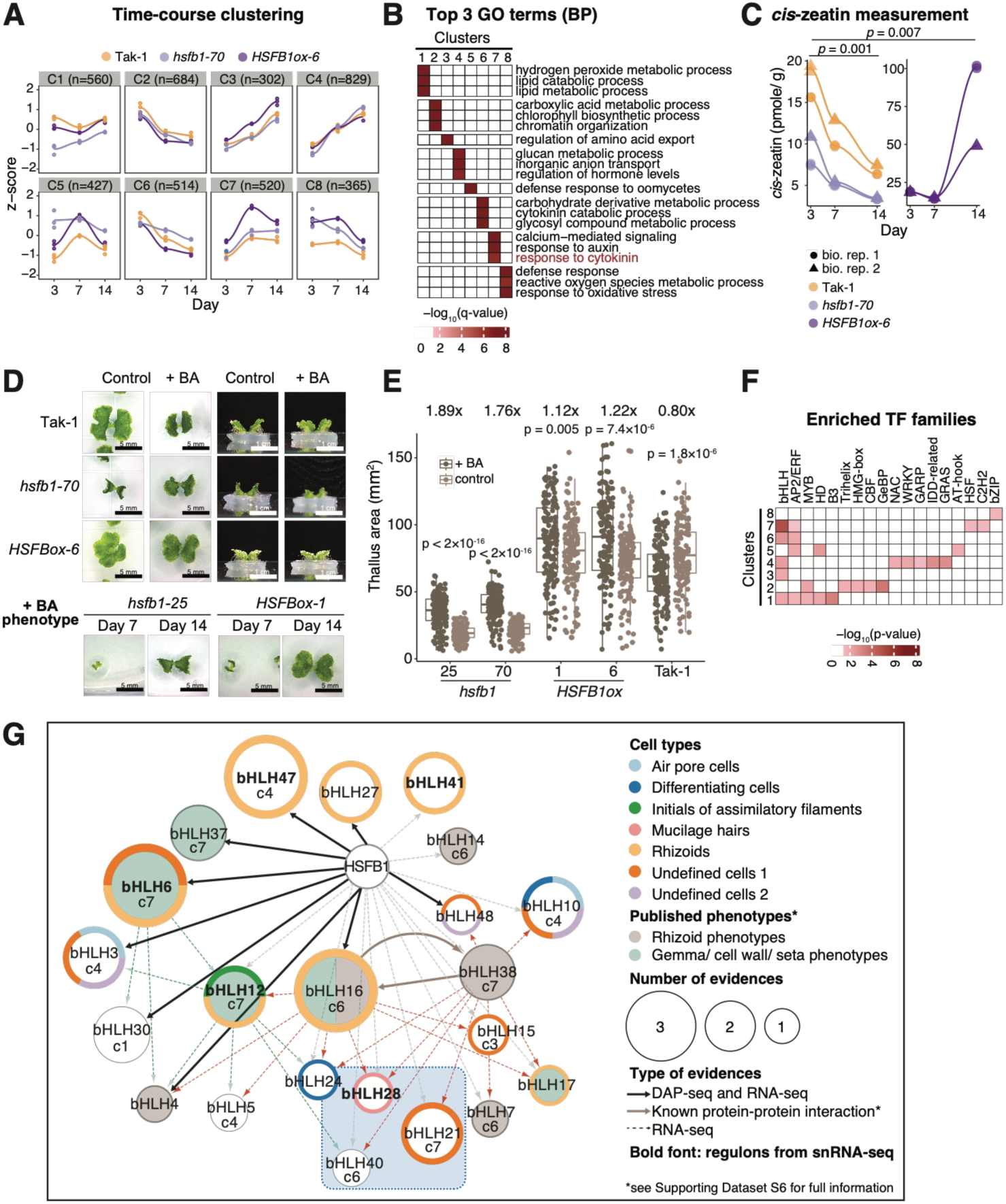
MpHSFB1 regulates cytokinin homeostasis during the early developmental stage of *Marchantia*. **A.** Hierarchical clustering of the time-course bulk RNA-seq dataset. The line represents the mean gene expression of each genotype group. **B.** Heatmap showing the top three significantly enriched GO (Biological Process, BP) terms from each cluster (q-value < 0.01). **C.** Line graphs showing cis-zeatin levels. Two biological replicates with two technical replicates each were performed for each genotype and condition. *p < 0.05 was determined by a pairwise Student’s t-test. **D.** Representative photos of cytokinin treatments in Tak-1, Mp*hsfb1*, and Mp*HSFB1ox* plants. Note that the photo for Tak-1 plants is from the same set as in Figure 6A (control condition), as the experiments were performed simultaneously. **E.** Boxplots showing thallus area for each genotype and condition. Dots represent individual plant data; p-values were determined by the Mann-Whitney U test with Bonferroni correction (*n* = 60 plants per line). Boxplots display all data points (dots), median values, first and third quartiles, and whiskers indicating minimum and maximum values. Fold change was calculated as the ratio of the mean thallus area between control and cytokinin treatments for each genotype. **F.** Heatmap showing the enrichment (hypergeometric distribution) of TF families in each cluster. Only TF families enriched in at least one cluster are shown. **G.** Integrated regulatory network of Mp*HSFB1*-mediated Mp*bHLH* regulation. The network integrates data from time-course RNA-seq clusters, public RNA-seq datasets, snRNA-seq datasets, regulons, DAP-seq, and known mutant phenotypes. Node size corresponds to the number of lines of supporting evidence (bulk RNA-seq, snRNA-seq, and DAP-seq). Solid lines represent physical DNA–protein or protein–protein binding interactions. Edge colors indicate whether the regulation is associated with a rhizoid- or gemma-phenotype-related bHLH gene. See Supporting Datasets S6 and S7 for detailed information.

We next sought to identify downstream transcriptional regulators that may connect MpHSFB1 to these cytokinin-associated developmental responses. The bHLH TF family was strongly enriched in the cytokinin-related clusters C6 and C7 (Figure 4F), while MpHSFB1-co-expressed MpbHLHs were also enriched in rhizoid cells based on snRNA-seq (Fisher’s exact test, *p* < 0.05; Supporting Dataset S2). We therefore focused on the regulatory relationship between MpHSFB1 and these MpbHLHs by integrating multiple genomic and transcriptomic datasets. DAP-seq identified several established developmental MpbHLHs, including MpbHLH4, 6, 16, and 37, as direct MpHSFB1-bound targets (Figure 4G, solid arrows)^23,24,27,28^. In parallel, DEGs identified in public Mp*bhlh* mutant datasets significantly overlapped with those altered in our Mp*hsfb1* mutant dataset, including MpBHLH7, 21, and 40 (Supporting Dataset S6 & S7). MpHSFB1 expression, however, remained unchanged in these Mp*bhlh* mutant backgrounds, supporting a regulatory direction from MpHSFB1 toward these bHLHs rather than the reverse. In addition, snRNA-seq identified MpbHLH28 as a key regulon component in mucilage hairs, which are structures involved in preventing water loss^29^ (Figures 3D & 4G). Based on these complementary analyses, we selected MpbHLH21, 28, and 40, which showed the strongest combined evidence across these analyses, for downstream functional characterization.

Luciferase reporter assays showed that MpHSFB1 was sufficient to repress the MpBHLH21 and MpBHLH28 promoters, but not the MpBHLH40 promoter (Supporting Figures S7 & S8, Supporting Dataset S6 & S7). These results indicate that MpHSFB1 regulates individual MpbHLHs through distinct mechanisms, with MpBHLH21 and MpBHLH28 showing direct evidence of transcriptional regulation by MpHSFB1, whereas the relationship with MpbHLH40 may involve additional regulatory components. Furthermore, we overexpressed MpHSFB1 in Mp*bhlh* knockout backgrounds to genetically test whether these bHLH factors function downstream of MpHSFB1. We compared developmental phenotypes of Tak-1, Mp*hsfb1-25*, Mp*HSFB1ox-1*, Mp*bhlh40-5*, Mp*bhlh28-18*, Mp*bhlh21-4*, and the corresponding double lines (Mp*bhlh40*/Mp*HSFB1ox*, Mp*bhlh28*/Mp*HSFB1ox*, and Mp*bhlh21*/Mp*HSFB1ox*). In all three genetic backgrounds, MpHSFB1 overexpression failed to override the respective Mp*bhlh* mutant phenotypes: the double lines retained the phenotypes of the corresponding Mp*bhlh* mutants in thallus size, solidity, and rhizoid-to-thallus ratio (Supporting Figure S9). The epistasis of the Mp*bhlh* mutations over MpHSFB1 overexpression provides genetic evidence that MpBHLH21, MpBHLH28, and MpBHLH40 function downstream of MpHSFB1 in developmental regulation.

We then asked whether these downstream MpbHLHs contribute to the developmental and cytokinin-related phenotypes associated with MpHSFB1. Mpb*hlh21* mutants displayed a significantly lower rhizoid-to-thallus ratio (∼0.67-fold), whereas Mp*bhlh40* mutants exhibited a higher ratio compared to Tak-1 (Figure 5A & 5B, ∼2.7-fold), resembling the rhizoid phenotype of Mp*hsfb1*. Beyond rhizoid development, Mp*bhlh28* mutants displayed deeply embedded gemmae within the gemma cups, whereas Mp*bhlh40* mutants failed to maintain dormancy (Supporting Figure S7B). Crucially, cytokinin levels were inversely correlated with rhizoid density (*ρ* = -0.67): Mp*bhlh21,* which produced fewer rhizoids, accumulated significantly higher cytokinin levels, whereas Mp*bhlh40,* which produced more rhizoids, showed significantly lower cytokinin levels (Figure 5C & 5D). Exogenous cytokinin partially rescued the Mp*bhlh40* mutant phenotype by restoring thallus area but had no detectable effect on Mp*bhlh21* mutants (Figure 5E & 5F), further linking MpBHLH40 to the low-cytokinin, high-rhizoid phenotype observed in the Mp*hsfb1* mutant.

**Figure 5.**
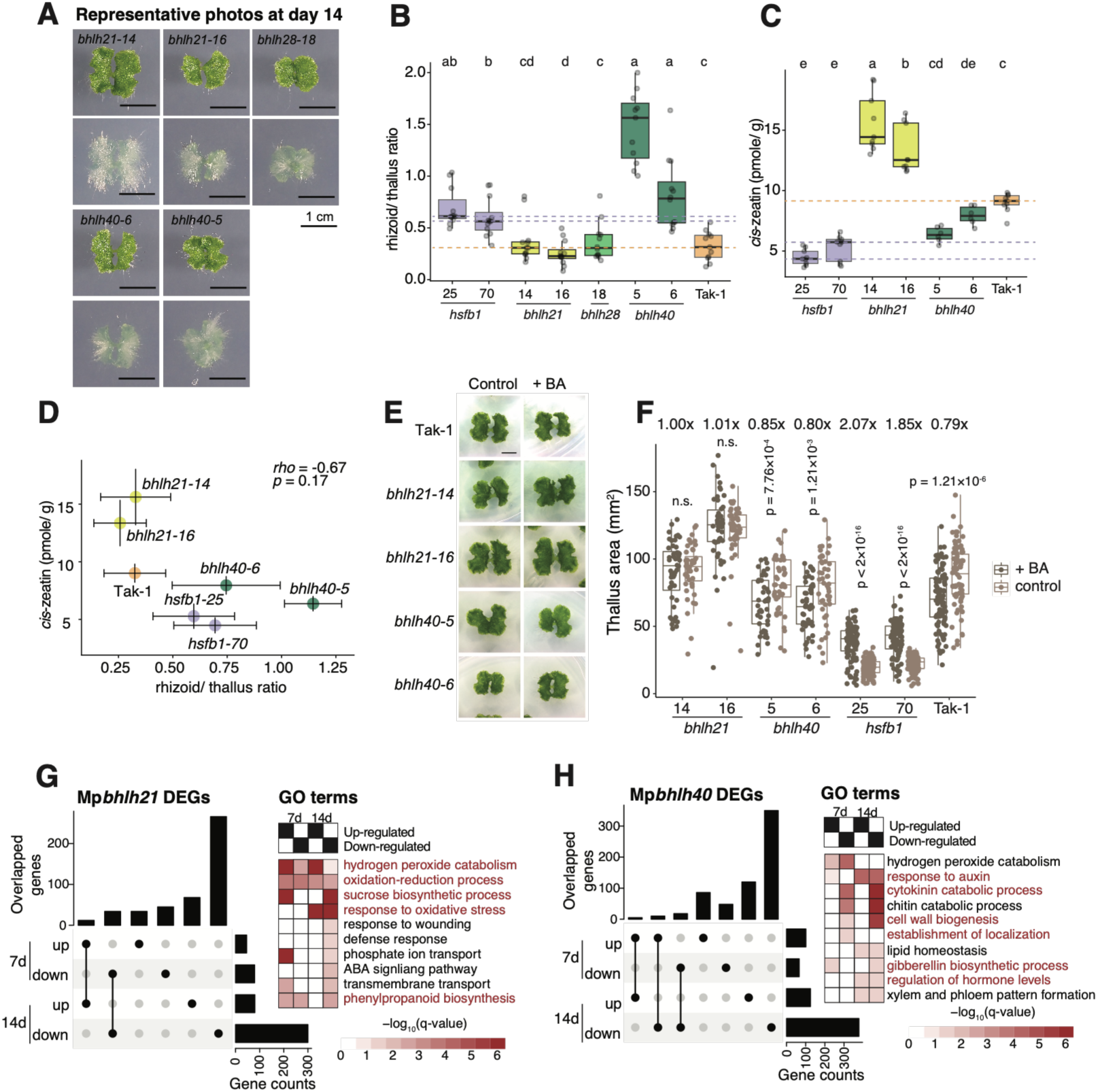
MpbHLHs regulate cytokinin homeostasis during the early developmental stage of *Marchantia*. **A.** Representative images of Mp*bhlh* knockout mutants at day 14. **B.** Boxplot showing the rhizoid-to-thallus ratio. P-values were determined by one-way ANOVA followed by Tukey HSD post hoc test (*n* = 10-12 plants per line). **C.** Boxplot showing the *cis*-zeatin levels. P-values were determined by one-way ANOVA followed by Tukey HSD post hoc test (*n* = 6-8 independent measurements). **D.** Scatter plot showing the Spearman correlation between the rhizoid/thallus ratio and cytokinin concentration. Each data point represents the mean value, with error bars indicating the standard error (S.E.). **E.** Representative photos of cytokinin treatments in Tak-1, Mpb*hlh21-14*, Mp*bhlh21-16*, Mp*bhlh40-5*, and Mp*bhlh40-6* plants. **F.** Boxplots showing thallus area for each genotype and condition. P-values were determined by the Mann-Whitney U test with Bonferroni correction (*n* = 30-60 plants per line). Fold change was calculated as the ratio of the mean thallus area between control and cytokinin treatments for each genotype. **G. & H.** UpSet plots and heatmaps showing genotype-specific DEGs of Mp*bhlh21* (**G**) and Mp*bhlh40* (**H**) mutants at days 7 and 14 relative to wild-type Tak-1 plants. For all panels, boxplots display individual data points (dots), median values, first and third quartiles, and whiskers indicating minimum and maximum values.

Consistent with these phenotypic differences, genotype-specific DEG analysis revealed distinct regulatory functions for these two MpbHLHs. Mp*bhlh40* DEGs were enriched in cytokinin responsiveness, pattern formation, and root development, whereas Mp*bhlh21* DEGs were associated with sucrose metabolism, abiotic stress, and ROS pathways (Figure 5G & 5H, Supporting Dataset S7). Integration with the Mp*hsfb1* and Mp*HSFB1ox* time-course profiles further identified shared target genes within clusters C1, C4, and C6 that were misregulated in both lines and enriched in ROS regulation, lipid metabolism, and hormone signaling (cytokinin and GA) (Supporting Dataset S7). Collectively, these regulatory, genetic, and physiological data support a model in which Mp*BHLH21* and Mp*BHLH40* function downstream of MpHSFB1 to coordinate distinct cytokinin-associated developmental and physiological programs.

### MpHSFB1 regulates prolonged HS tolerance via modulating ROS response

Regulating ROS homeostasis is essential for thermotolerance under prolonged HS^30^. Under prolonged HS, Mp*HSFB1ox* lines exhibited accelerated pigmentation and significantly reduced thallus area compared to Tak-1 and Mp*hsfb1* plants (Figure 6A, Supporting Figure S10). While ROS levels decreased significantly in Tak-1 and Mp*hsfb1* lines post-HS (*p* = 1.8 × 10⁻¹³ and 0.018, respectively), they remained stable in Mp*HSFB1ox* lines, suggesting that overexpression prevents proper ROS suppression and triggers oxidative stress under prolonged HS (Figure 6C, Supporting Figures S10C & S10D, Supporting Dataset S1). RNA-seq identified hundreds of genotype-specific DEGs, including 35 with opposite expression patterns between genotypes (Figure 6D, Supporting Figure S10B, Supporting Dataset S8). These targets were enriched in ROS responses, oxidative stress, and pigment biosynthesis (Figure 6E), supporting the hypothesis that MpHSFB1 plays a pivotal role in maintaining ROS homeostasis during temperature adaptation.

**Figure 6.**
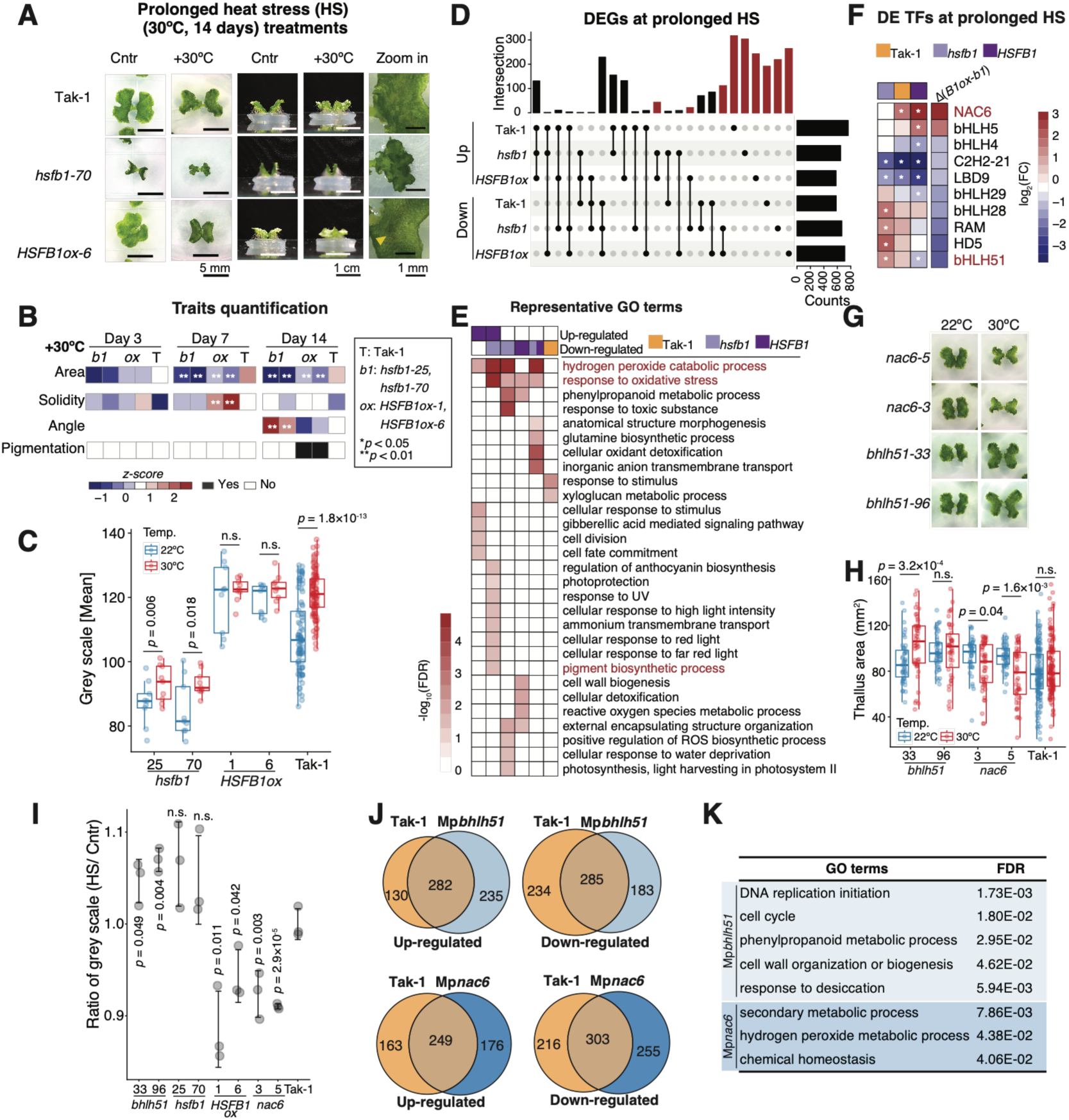
MpHSFB1 regulates prolonged HS tolerance via the ROS response. **A.** Representative top-view and side-view images after prolonged HS treatment at day 14. **B.** Heatmaps showing quantitative analyses of developmental traits. Z-scores were calculated across developmental stages and genotypes. p-values were determined using Student’s t-test between mutants and the wild type. Two independent experiments were performed with *n* = 30–60 per time point. Raw measurements and complete statistical results are provided in Supporting Dataset S1. **C.** Boxplots showing DAB staining intensities in each genotype and condition. The y-axis denotes grayscale pixel values (0 [black]–255 [white]). P-values were determined by the Mann-Whitney U test with Bonferroni correction (*n* = 12–60 plants per line). **D.** UpSet plot showing up- and down-regulated DEGs under prolonged HS. The red bars denote Mp*hsfb1*- or Mp*HSFB1ox*-specific genes subjected to further analysis. **E.** Heatmap showing representative enriched GO (BP) terms (FDR < 0.05) from the groups in (D). **F.** Heatmap showing log₂FC values of selected TFs (|Δ(*B1ox* - *b1*)| ≥ 1) across genotypes under prolonged HS, highlighting expression differences between Mp*HSFB1ox* lines and Mp*hsfb1* mutants. Asterisks denote DE TFs, and red labels indicate TFs selected for further validation. **G.** Representative images of Mp*bhlh51* and Mp*nac6* knockout mutants at day 14 under prolonged HS. **H.** Boxplots showing thallus area for each genotype and condition. P-values were determined by the Mann-Whitney U test with Bonferroni correction (*n* = 60 plants per line). **I.** Dot plot showing the grayscale ratio under prolonged HS relative to control conditions for each genotype. Error bars indicate the standard error (S.E.). A lower ratio indicates a higher ROS level. *p < 0.05 was determined by a pairwise Student’s t-test. **J.** Venn diagram showing the number of genotype-specific DEGs in *Mpbhlh51* and *Mpnac6* mutants under prolonged HS conditions. **K.** Significantly enriched GO BP terms corresponding to the DEGs identified in (J). For all panels, boxplots display individual data points (dots), median values, first and third quartiles, and whiskers indicating minimum and maximum values.

Expression profiling revealed that Mp*BHLH51* and Mp*NAC6* (MpJINGASA^31^) exhibited the most pronounced differences in expression between Mp*HSFB1ox* lines and Mp*hsfb1* mutants (Figure 6F, Supporting Figure S10E). Functional analyses of their respective mutants revealed divergent regulatory roles, where Mp*bhlh51* mutants developed significantly larger thalli (Figures 6G & 6H) and accumulated less ROS under stress (Figure 6I, Supporting Figure S10F), consistent with an enrichment of their DEGs in generalized stress adaptations such as ROS detoxification, cell wall reorganization, and phenylpropanoid metabolism (Figures 6J & 6K, Supporting Dataset S8). Conversely, Mp*nac6* mutants displayed severe hypersensitivity to prolonged HS, characterized by restricted growth and elevated ROS accumulation, which aligns with the enriched function that was specialized in redox homeostasis and osmotic stress (Figure 6G-6K). Together, these contrasting downstream cascades demonstrate that MpHSFB1 acts as a central coordinator of thermal adaptation by fine-tuning distinct, specialized regulatory modules.

To test the regulatory specificity of downstream pathways, we performed cross-phenotypic evaluations of the development- and stress-related mutants. When challenged with prolonged HS, the development-associated Mp*bhlh21* and Mp*bhlh40* mutants maintained a thallus area and ROS ratio indistinguishable from wild-type Tak-1 plants, indicating they did not participate in prolonged HS (Supporting Figure S11A & S11B). Reciprocally, under control conditions, the stress-responsive Mp*bhlh51* and Mp*nac6* mutants, as well as Mp*bhlh51*/Mp*HSFB1ox* lines, exhibited no significant abnormalities in their rhizoid-to-thallus ratio or endogenous cytokinin concentrations (Supporting Figure 9, Supporting Figure S11C - S11E). Together, these results support distinct functional roles for these downstream regulators, with MpBHLH21 and MpBHLH40 primarily associated with developmental and cytokinin-related processes, and MpBHLH51 and MpNAC6 with prolonged HS responses.

### MpHSFB1 and its orthologs coordinate tissue- and cell type-specific transcriptional programs in plants

Co-expression networks across species revealed that distinct HSFBs serve as regulatory hubs within different cell types (Supporting Dataset S9). For example, HSFB2a was top-ranked in *Arabidopsis* meristem procambium cells and the rice epidermis, while HSFB1 acted as a hub in tomato and maize root cap cells (Supporting Dataset S9). Based on GO term similarity, the functions of the genes associated with each HSFB module were grouped into seven clusters (Supporting Figure S12A, Supporting Dataset S9). Focusing on these functions, MpHSFB1 showed a broad, even distribution across GO groups, whereas angiosperm HSFB modules displayed narrower, more specialized functional distributions. Specifically, genes associated with AtHSFB2a, SlHSFB1, and SlHSFB3a exhibited a significantly higher proportion of stress-related functions and a depletion of development-associated functions (*p* < 0.05 compared to MpHSFB1) (Supporting Figure S12B). These data suggest that the ancestral MpHSFB1 regulates a broad, evenly distributed array of functional target pathways, whereas angiosperm HSFBs have shifted toward more specialized functional target profiles.

We next investigated the conservation of HSFB regulatory logic at the cellular level. Since the expression patterns of a TF’s conserved target genes allow us to chart TF activity across species^32^, we performed swapDAP-seq to assess the binding capacity of MpHSFB1 on the *Arabidopsis* genome (Figure 7A, Supporting Figure S13A, Supporting Dataset S10). By profiling MpHSFB1 binding to both Marchantia and *Arabidopsis* genomic DNA, we identified 535 target genes (392 orthogroups) and 2,302 target genes (2,082 orthogroups), respectively (Figure 7B). Only 63 overlapping orthogroups (2.6%) were shared between the two species, which were strictly enriched for core heat stress response (FDR = 1.85 × 10^-6^) and ROS response (FDR = 0.011). Conversely, the closest *Arabidopsis* homolog, AtHSFB4, bound far fewer genomic sites, yielding no overlapping orthogroups between genomes (Figure 7C)^33^. Furthermore, its *Arabidopsis*-specific targets were tightly restricted to specific localized functions, such as malate transport (FDR = 5.93 × 10^-3^) and cell division (FDR = 0.013) (Figure 7D). Thus, genomic targeting has diverged within the HSFB clade, where MpHSFB1 retains broad developmental and stress-responsive binding potential, whereas tested angiosperm HSFBs display significantly narrower profiles.

**Figure 7.**
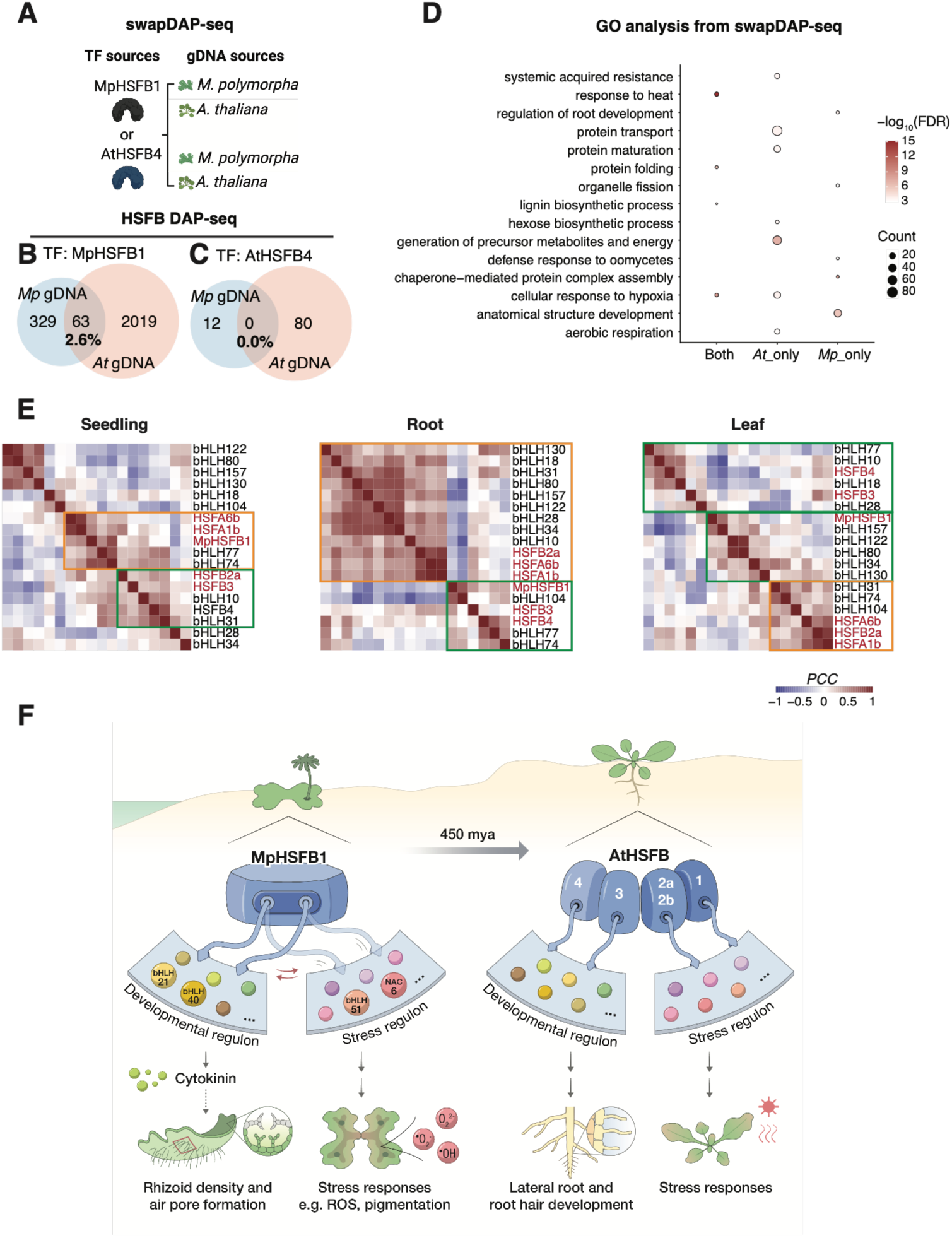
MpHSFB1 and its orthologs coordinate tissue- and cell type-specific transcriptional programs across land plants. **A.** Schematic illustrating the experimental design for swapDAP-seq. **B & C.** Venn diagrams showing the overlapping orthogroups identified by swapDAP-seq using MpHSFB1 (**B**) or AtHSFB4 (**C**) bound to the Marchantia and Arabidopsis genomes. **D.** Heatmap showing the enriched GO terms identified from genes bound exclusively in Arabidopsis, exclusively in Marchantia, or in both species from (**B**.). **E.** Heatmap showing the similarity of TF regulation across four Arabidopsis tissues, calculated using the Pearson correlation coefficient. **F.** Proposed role of MpHSFB1 in land plant adaptation. The model illustrates how MpHSFB1 coordinates distinct transcriptional programs for development and stress resilience. In *Marchantia*, MpHSFB1 cooperates with specific bHLH partners to regulate cytokinin homeostasis during early development and ROS responses under prolonged heat stress. This regulatory flexibility highlights the co-option of HSFB1 networks in orchestrating tissue-specific adaptations across land plants.

Finally, we validated these regulatory shifts using publicly available snRNA-seq and scRNA-seq datasets alongside our own *Marchantia* snRNA-seq data (Supporting Figure S14, Supporting Dataset S10). While Arabidopsis HSFB regulon activity is broadly enriched in undifferentiated meristematic cells across seedling, leaf, and root tissues, MpHSFB1 activity is strongly associated with specialized, differentiated cell types like air pores and oil body cells (Figure 2). Notably, MpHSFB1 shared regulatory features with both stress-responsive cell types (resembling AtHSFB2a and AtHSFAs) and developing cell types (resembling AtHSFB3 and AtHSFB4) (Supporting Figure S14). This cell-type-dependent versatility is associated with distinct HSFB–bHLH co-regulatory arrangements (Figure 7E). In Arabidopsis roots, AtHSFB3 and AtHSFB4 correlated positively with cell division-regulating TFs (AtbHLH074, AtbHLH77, and AtbHLH104). Similarly, MpHSFB1 exhibited dual regulatory associations: it correlated strongly with stress-related bHLHs and HSFBs in seedling cells, but with development-related TFs in roots and leaves (Figure 7E). These findings suggest a model where cell-type-specific HSFB–bHLH interactions potentially underlie distinct adaptive traits in *Marchantia* and *Arabidopsis* (Figure 7F).

## Discussion

Overcoming terrestrial challenges was a pivotal step in land plant evolution, coinciding with the diversification and expansion of the HSFB and bHLH transcription factor families. These events enabled the regulatory innovations required for life on land. In this study, we demonstrate that MpHSFB1 functions as an ancestral regulatory hub that coordinates distinct bHLH and NAC transcription factor modules to orchestrate developmental and stress-responsive programs across different cell types (Figures 2-7). By integrating cytokinin signaling, MpHSFB1 modulates bHLH21 and bHLH40 to fine-tune the spatiotemporal dynamics of rhizoid development, while independently coordinating bHLH51 and NAC6 to mediate adaptive tolerance under prolonged heat stress. Such modularization illustrates how conserved regulatory networks were progressively co-opted and rewired to generate new adaptive traits during land plant evolution. Collectively, our findings suggest that ancestral HSFB transcriptional regulons were repurposed through regulatory rewiring to facilitate the specialized cellular functions during land plant evolution.

The transition of plants to land required the emergence of new cell types to form tissues and organs, accompanied by the evolution of sophisticated tissue-to-tissue communication systems. The evolution of GRNs involving cytokinin signaling likely facilitated the coordinated regulation of stress acclimation and developmental processes in land plants (Figures 3, 5, and 7)^34^, including the plastic development of root or rhizoid systems and the accumulation of protective pigments such as anthocyanins or auronidins^35^. Indeed, endogenous cytokinin concentrations were misregulated in Mp*hsfb1* and Mp*bhlh* mutants, and exogenous cytokinin application partially rescued their developmental phenotypes (Figure 5), supporting the hormone’s role in integrating the molecular cues underlying these adaptive traits. Furthermore, HS is known to reduce cytokinin content, whereas exogenous cytokinin generally enhances HS resistance^36-38^. Cytokinin and HS responses cross-talk by regulating HSPs and photosynthesis-related proteins, thereby influencing stomatal movement^38^. Given the misregulation of cytokinin signaling and cytokinin-responsive genes in Mp*hsfb1* mutants, it is likely that the integration of HSFB into stress- and hormone-responsive pathways functioned as a critical adaptive signaling mechanism during plant evolution (Figure 7).

We propose that MpHSFB1 represents an evolutionarily ancient HSFB that originally possessed multimodal functions, which later diverged through subfunctionalization within the angiosperm lineage. In *Marchantia*, MpHSFB1 regulates both developmental and stress-related processes (Figures 5 and 6). By contrast, the *Arabidopsis* genome encodes five HSFB genes with specialized roles: AtHSFB1, AtHSFB2a, and AtHSFB2b are primarily involved in stress regulation, whereas AtHSFB3 and AtHSFB4 function mainly in developmental control. For example, AtHSFB4 (SCHIZORIZA) regulates root cell-fate segregation, while the closest homolog of AtHSFB3 in black cottonwood controls xylem cell-wall biosynthesis^13,39^. Interestingly, despite sharing nearly identical binding motifs, AtHSFB3 and AtHSFB4 targeted fewer genes in DAP-seq datasets than AtHSFAs or AtHSFB2a. A similar trend was observed when comparing MpHSFA1 and MpHSFB1. In our cross-species comparison, we found that MpHSFB1 is capable of binding to *Arabidopsis* genomic DNA, whereas AtHSFB4 cannot bind to the *Marchantia* genome. In contrast, HSFA1 proteins exhibit conserved binding capacities across species boundaries (Figure 7, Supporting Figure S13). Consistent with this, the HSFB lineage underwent strong positive selection in bryophytes relative to the ancestral HSFs found in algae, likely driven by the drastic environmental shifts associated with land plant adaptation^10^. These results indicate that sequence divergence within the HSFB DNA-binding domain (DBD) was a key determinant of the functional diversification and specialization observed during plant evolution.

While direct protein–protein interactions between MpHSFB1 and the MpbHLHs were undetected (Supporting Figure S7), DAP-seq revealed that the identified motifs were generally longer in Marchantia (Figure 5)*^40,41^*. Because many TF families form higher-order complexes to recognize longer motifs and increase DNA-binding specificity, these longer motifs suggest the HSFB family may form similar complexes to enhance targeting specificity and enable modular control^42-44^. Coupled with evidence indicating that HSFBs cooperate with distinct bHLH TFs in a cell-type-specific manner (Figure 7), these findings illustrate how TFs can be co-opted to generate new, cell-type-specific regulatory functions.

Like other eukaryotic TFs, bHLH factors form distinct regulatory complexes to govern diverse developmental and stress responses. For example, the *Arabidopsis* homologs of MpBHLH40 (AtbHLH037 and AtbHLH088) control reproductive development and photomorphogenesis^45^, while MpBHLH21 and MpBHLH28 homologs regulate cell elongation and root hair growth, indicating this functional diversity had already emerged in bryophytes. This regulatory versatility may be driven by several mechanisms: association with ACT-like domains, post-translational control, or the cell-type-specific modulation of hormone pathways like cytokinin signaling^46^. Together, these features underscore how bHLH TFs drove the diversification of plant developmental and stress-responsive programs. In conclusion, these results suggest how ancestral HSFB-mediated regulatory modules evolved to facilitate cell-type specificity and functional diversity in land plants.

## Material and Methods

### Phylogenetic tree construction

Full-length protein sequences were retrieved from the Tapscan database^47^, JGI PhycoCosm^48^, and recent genome releases^49^. Our dataset encompasses key evolutionary lineages, ranging from streptophyte algae (*Klebsormidium nitens [v1.0]*, *Mesotaenium endlicherianum [SAG12.97]*, *Zygnema circumcarinatum [Zygcir6981b_2]*) to bryophytes (*Anthoceros agrestis [v1.0]*, *Marchantia polymorpha [v6.1r2]*), tracheophytes (*Selaginella moellendorffii [v1.0]*, *Azolla filiculoides [v1.2]*, *Salvinia cucullata [v2]*, *Picea abies [v1]*, *Ginkgo biloba [v1]*, *Oryza sativa [MSU v7.0]*, *Zea mays [B73_v4]*, *Arabidopsis thaliana [Araport11]*, *Solanum lycopersicum [ITAG4.1]*), with *Chlorophyta* and *Rhodophyta* species serving as outgroups. Conserved DBDs were identified using the NCBI Conserved Domain Database. We performed multiple sequence alignment via MAFFT (v7.520) using the L-INS-i algorithm (--maxiterate 1000 --localpair) and reconstructed the maximum-likelihood phylogenetic tree using IQ-TREE (v2.2.2.6). The optimal amino acid substitution model was selected via ModelFinder Plus based on the Bayesian Information Criterion, with branch support evaluated using 1,000 Ultrafast Bootstrap replicates.

### Growth condition and morphological characterization

*Marchantia polymorpha* wild type (accession Tak1), MpHSFB1 overexpression lines (Mp*HSFB1ox*), MpHSFB1 knockout lines (Mp*hsfb1*), and TF candidate knockout mutants were grown on 1/2 × Gamborg B5 medium (pH 5.5) with 1% agar. Plants were grown under continuous light at either 22 °C or 30 °C, or with and without the treatment of 10 μM 6-Benzylaminopurine (BA, Sigma). Morphological characterization was performed in two phases: developmental and treatment. For the developmental phase, plants were cultured at 22 °C, and images were captured on days 3, 7, and 14. For the treatment phase, plants were cultured at either 22 °C or 30 °C, with or without BA treatment, and images were captured on day 14. All images were obtained using a Zeiss Lumar V12 microscope, while other images were captured using a digital camera and further analyzed using Fiji ImageJ software.

### Phenotypic analysis

#### Thallus area and shape analysis

The scale of each image was calibrated by drawing a straight line along the scale bar (Set Scale). Measurement parameters, including area, shape descriptors, and area fraction, were selected. Each image was then split into red, green, and blue channels (Split Channels), and subsequent analyses were performed on the blue channel. To reduce image noise, a Gaussian blur filter was applied (Gaussian Blur). Images were automatically thresholded to distinguish the thallus from the background. Non-thallus regions were cropped to retain only the area of interest, and the image was inverted to prepare for particle analysis. Thallus area and shape characteristics were quantified using “Analyze Particles,” and the corresponding values were recorded. The convex hull of each thallus area was generated (Convex Hull), and measurements were repeated to obtain convex hull area values.

#### Rhizoid area analysis

For rhizoid quantification, the scale was calibrated, and an area of interest was selected. Each image was split into red, green, and blue channels, and either the red or blue channel was used, depending on which provided better contrast between rhizoids and the background. Thresholding and selection of rhizoid areas were performed manually. Non-rhizoid regions were cropped to retain only the rhizoid area, and the image was inverted prior to measurement.

#### Angle measurement

Side-view images were used to measure plant shoot angles. Each image was cropped to include only the green portion of the plant. The Angle tool in ImageJ was used for measurement. For the left-side angle, three points were selected sequentially: (1) the top-left corner of the plant, (2) the midpoint of the plant, and (3) a point forming a horizontal line with the midpoint on the left side, after which the angle was measured. For the right-side angle, the midpoint was maintained, while the first point was moved to the top-right corner and the third point to the right side, again forming a horizontal line with the midpoint. Measurements were repeated as described above.

### Overexpression and knockout lines in *Marchantia*

The MpHSFB1 overexpression construct was generated using Mol-Clo–based Golden Gate assembly. The coding sequence (CDS) of MpHSFB1 (synthesized by OmicBioscience) was fused to an N-terminal 3×Flag tag and a C-terminal mCherry and cloned under the control of the CaMV35S promoter with a NOS terminator to generate a Level 1 (Lv1) plasmid. This Lv1 plasmid was subsequently assembled with a second Lv1 plasmid containing a hygromycin resistance cassette driven by the MpEF1α promoter, yielding the Level 2 (Lv2) plasmid used for Agrobacterium-mediated transformation^9^. To complement Mp*bhlh* mutants with MpHSFB1, the CmmT control construct (CaMV35S:mCherry:YFP:miniTurbo:NOS) and the CBmT construct expressing MpHSFB1 (CaMV35S:mCherry:MpHSFB1:miniTurbo:NOS) were generated via Golden Gate assembly. Successful transformants were screened on 1/2 × B5 medium plates containing 100 µg/mL cefotaxime or 10 mg/L hygromycin. Two independent overexpression lines at the T2 generation were selected based on transgene expression levels determined by qPCR.

The Mp*hsfb1* mutant was generated in our previous publication^9^. Knockout mutants of MpbHLH21, MpbHLH28, MpbHLH40, and MpbHLH51 were produced using the CRISPR/Cas9 (clustered regularly interspaced short palindromic repeats/CRISPR-associated protein 9) genome-editing system. Two guide RNAs (gRNAs) targeting sequences flanking a short DNA fragment within each locus were designed and cloned into either the pMpGE010 (Hygromycin resistance) or pMpGE011 (chlorsulfuron resistance) expression vector. Successful transformants were screened on 1/2 × B5 medium plates containing 100 µg/mL cefotaxime and 0.5 µM chlorosulfuron or 10 mg/L hygromycin. The mutation sites of two independent knockout lines at the T2 generation were confirmed via PCR and sequencing. Gene locus identifiers and primers used for sequencing and cloning are listed in Supporting dataset S11.

### Detection of reactive oxygen species (ROS)

The histochemistry staining was followed by a previous study^16^. *Marchantia* thalli were harvested from 14-day-old Tak-1, Mp*HSFB1ox* lines, Mp*hsfb1* lines, and TF candidate knockout mutants and stained with 3,3ʹ-Diaminobenzidine (DAB). The 1 mg ml−1 DAB staining solution was directly applied on the plate, followed by 5 min vacuum and shake for 1 h, at 50rpm in the dark incubation. The DAB staining solution was replaced with distilled water, subsequently with 75% ethanol to remove the chlorophyll. Plant images were taken and analyzed by Fiji ImageJ software.

### Endogenous cytokinin detection

Thalli from 3-, 7-, and 14-day-old Tak-1, Mp*HSFB1ox*, and Mp*hsfb1* lines were ground using a pestle and resuspended in Bieleski buffer (60% methanol, 25% CHCl_3_, 10%HCOOH, and 5% H2O). The mixture was sonicated on ice for 30 min, followed by centrifugation at 13,000 rpm for 10 minutes at 4°C. The supernatant was transferred into a new 1.5 mL tube and dried using a vacuum concentrator at 4 °C. UPLC-MS/MS analysis was performed by ABRC Metabolomics Core. Deuterium-labeled trans-zeatin (tZ-d5) was used as an internal control. Absolute quantification of cytokinins was calculated using the following formula: [cZ_(area)_/d5-tZ_(area)_ x_0.5_ng_(d5-tZ)_]÷FW_(mg)_= ng/mg.

### Oil Body Staining

For oil body observation, gemmae were collected from Tak-1 plants, Mp*HSFB1ox* lines, and Mp*hsfb1* mutants. The fluorophore BODIPY 493/503 (4,4-difluoro-1,3,5,7,8-pentamethyl-4-bora-3a,4a-diaza-s-indacene; Thermo Fisher Scientific) was dissolved in water to prepare a 0.5 mg/mL stock solution, which was stored at -20 °C. Prior to imaging, the tissues were incubated in a 0.05 mg/mL BODIPY working solution for 10 min, followed by two washes with ddH_2_O. Fluorescence was visualized using a Zeiss LSM 880 confocal microscope with a 488 nm excitation argon laser. Finally, the number of oil bodies per gemma was quantified manually.

### Protein-protein interaction (PPI) assay using RUBY-based reporter system

The MpHSFB CDS was fused to the N-terminal GAL4 DNA-binding domain (GAL4) under the control of a double CaMV35S promoter. Candidate TF CDSs were synthesized (by Twist Bioscience), cloned in-frame with the C-terminal GAL4 activation domain (p65), and driven by the double CaMV35S promoter. The three constructs were introduced into Agrobacterium tumefaciens by electroporation. Agroinfiltration was performed as previously described with minor modifications^50^. Briefly, transformed Agrobacterium strains were cultured in 5 mL LB broth supplemented with 250 µg/mL carbenicillin, 10 mM MES (pH 5.7), and 20 μM acetosyringone at 28 °C for 2 days. Cells were collected by centrifugation and resuspended in infiltration buffer (10 mM MgCl₂, 10 mM MES, pH 5.7, and 200 μM acetosyringone) to an OD₆₀₀ of 1.0, then incubated at room temperature for at least 1 h. Equal volumes of the three Agrobacterium suspensions were mixed (1:1:1) and infiltrated into fully expanded leaves of 4-week-old *Nicotiana benthamiana* plants using a needleless syringe. Betalain pigment accumulation was monitored 2 days post-infiltration.

### Generation and analysis of the DAP-seq dataset

#### Plasmid construction

The coding sequences (CDS) of MpHSFB1 were synthesized by Twist Bioscience, and the CDS of Arabidopsis HSFB4 (AtHSFB4) was amplified from *Arabidopsis* cDNA and subcloned to Lv0 acceptor pAGM9121. The CDSs were cloned downstream of the SP6/T7 promoter with an N-terminal HaloTag and a T7 terminator to generate level 1 (Lv1) expression vectors. Each CDS was first subcloned into pAGM9121 as a level 0 module using BbsI, and subsequently assembled into the pICH47732 Lv1 vector via Golden Gate cloning, yielding an SP6/T7 promoter–driven construct encoding a HaloTag-fused CDS followed by a T7 terminator. Recombinant proteins were expressed in vitro using the TNT SP6/T7 High-Yield Wheat Germ Protein Expression System (Promega) according to the manufacturer’s instructions. Protein expression and purity were verified by SDS–PAGE.

#### DNA Fragmentation and Preparation

Approximately 100 mg of *Marchantia* gemmaling (14 days old) were harvested and ground in liquid nitrogen. Genomic DNA was extracted using the CTAB-phenol/chloroform method and sheared mechanically to an average size of 300–500 bp. DNA fragments were end-repaired, A-tailed, and ligated to indexed sequencing adapters for multiplexing. Library preparation, including fragmentation and adapter ligation, was performed by the Genomic Technology Core Facility at IPMB.

#### Formation of Protein-DNA Complexes

*In vitro*–translated proteins were immobilized on HaloTag magnetic beads (Promega) by one hour incubation and washed three times with PBS containing 0.005% NP-40 to remove unbound proteins. The purified TF–bead complexes were then incubated with 100 ng of adapter-ligated genomic DNA and 10 μg of Sheared Salmon Sperm DNA (Thermo Fisher) in binding buffer at room temperature for 1 hour to allow specific protein–DNA interactions. Following incubation, beads were magnetically separated and washed three times to eliminate nonspecific DNA. Bound DNA was eluted by heating at 98°C for 10 minutes.

#### DAP-seq analysis

Raw sequencing reads were trimmed using Trimmomatic (v0.39)^51^ to remove adapter sequences and low-quality bases, specifically employing a SLIDINGWINDOW:4:15 quality filter and retaining reads with a minimum length of 36 bp. Trimmed reads were aligned to the *Marchantia* reference genome (v6.1r2) or *Arabidopsis* reference genome (TAIR 10) using Bowtie2 (v2.4.4)^52^ in end-to-end mode with default parameters. Significant binding regions were identified using MACS2 (v2.2.7.1)^53^ against a corresponding control sample, employing default settings. Peaks were retained if the false discovery rate (FDR) with Benjamini-Hochberg correction was < 0.05. MpHSFB1 or AtHSFB4 peak summits were assigned to target genes using the criterion as follows: a peak was associated with a gene if its summit overlapped the proximal promoter region (-1,000 bp to +500 bp relative to the annotated TSS). A Halo-tagged GFP was used for negative control. For motif analysis, a 100 bp DNA sequence centered on each peak summit (±50 bp) was extracted using BEDTools getfasta^54^. These sequences were analyzed for enriched motifs using MEME^55^, allowing for motif widths ranging from 6 to 20 base pairs with default background correction.

### Luciferase assay

*Plasmid construction:* Dual-luciferase reporter constructs were assembled via Golden Gate cloning, with each binary vector containing three transcriptional units at defined positions. Specifically, position P1 carried either a CaMV 35S promoter-driven MpHSFB1-6×HA effector cassette or an empty vector control; position P2 contained a firefly luciferase (LUC2) reporter cassette driven by the candidate bHLH promoter (or a 35S minimal promoter fused to the bHLH promoter); and position P3 contained a CaMV 35S promoter-driven Renilla luciferase cassette to serve as an internal normalization control. All bHLH promoters were amplified from Tak-1 genomic DNA using the specific primers listed in Supporting Dataset S11.

#### Agrobacterium-mediated transient expression in Nicotiana benthamiana

Binary vectors were electroporated into *Agrobacterium tumefaciens* strain GV3101. Individual colonies were cultured for 2 days in selective LB medium at 28°C with shaking, pelleted by centrifugation, and resuspended in infiltration buffer (10 mM MgCl₂, 10 mM MES, pH 5.6, and 200 μM acetosyringone) to an OD₆₀₀ of 0.1. Suspensions were incubated at room temperature for at least 30 minutes before being infiltrated into the abaxial side of 4-week-old *Nicotiana benthamiana* leaves using a needleless syringe. For each construct, infiltrations were performed on at least three independent leaves across three separate plants. At 48 hours post-infiltration, leaf tissue was harvested for luciferase measurements.

#### Dual-Glo luciferase assay

Infiltrated *N. benthamiana* leaf tissue was collected from each infiltration area, immediately frozen in liquid nitrogen, and homogenized in 150 μL Passive Lysis Buffer (E1941, Promega). Lysates were clarified by centrifugation at 12,000 *g*1 5–10 min at 4°C, and the supernatants were transferred to a white 96-well plate.Firefly and Renilla luciferase activities were measured using the Dual-Glo® Luciferase Assay System (E2920, Promega) according to the manufacturer’s instructions. Briefly, 25 μL of clarified leaf extract was mixed with an equal volume of Dual-Glo® Luciferase Reagent. After incubation for 10 min at room temperature, firefly luminescence was measured using a plate reader (SpectraMax i3X, Molecular Devices). Subsequently, an equal volume of Dual-Glo® Stop & Glo® Reagent was added to each well. After incubation for 10 min at room temperature, Renilla luminescence was measured. Two technical replicates were averaged before statistical analysis. Each independently infiltrated leaf area was treated as one biological replicate.

### Generation of single-nuclei RNA-seq (snRNA-seq)

#### Nuclei isolation and snRNA-seq procedure

A total of 2.0 grams of 7-day-old *Marchantia* plants were chopped gently using an ice-cold razor blade in 2 mL of ice-cold modified Galbraith buffer (45 mM MgCl2, 20 mM MOPS, 30 mM sodium citrate, 0.5% Triton X-100, 0.2 U/mL RNaseOUT™ Recombinant Ribonuclease Inhibitor, and 2 mM/mL DTT, pH 7.0) for 4 minutes in a chilled petri dish, followed by incubation in a 4°C fridge for 10 minutes. The lysate was subsequently filtered sequentially through 70 µm and 40 µm cell strainers and transferred into an ice-cold 50 mL Falcon tube. Nuclei were pelleted at 500 × *g* at 4°C for 5 minutes, after which the supernatant was carefully removed. The nuclei pellet was gently resuspended in 1 mL of modified LB01 buffer (15 mM Tris pH 7.5, 0.5 mM Spermine, 10 mM KCl, 20 mM NaCl, 0.1 U/mL RNaseOUT™ Recombinant Ribonuclease Inhibitor, and 2 mM/mL DTT) and filter through 30 µm cell strainer into a 5 mL (12 × 75 mm) round-bottom tube followed by 2 µg/ml DAPI staining.

Flow cytometry was performed using a Beckman CytoFLEX SRT equipped with a 100-µm nozzle and operated at a low flow rate (psi < 20). The photomultiplier settings were as follows: Forward Scatter (FSC) at 25 V, Side Scatter (SSC) at 25 V, and V450 (DAPI channel) at 100 V. Sorting gates were empirically defined using FSC and SSC to exclude debris and potential doublets. To establish baseline fluorescence, FACS analysis was conducted on unstained control samples before sorting DAPI-stained nuclei. For nuclei isolation, sort gates were set to specifically collect DAPI-positive nuclei. A total of 400,000 sorted nuclei were collected into 200 µL of 1X PBS + 1% BSA (0.4 U/mL RNaseOUT™ Recombinant Ribonuclease Inhibitor) in 1.5 mL Eppendorf tubes within 30 minutes. The nuclei were then pelleted by centrifugation at 500 × *g* for 10 minutes at 4°C. The supernatant was carefully removed, leaving approximately 80 µL. The quality and yield of nuclei were checked under EVOS M7000 with C-Chip using the DAPI channel. About 16,000-19,000 nuclei of each sample were loaded into the 10X Genomics chip for cell barcoding. The libraries were prepared following the manufacturer’s instructions with a 10X Chromium Single Cell 3’ Solution v3.1 kit. The paired-end libraries were sequenced with the Illumina NextSeq2000 instrument at the NGS Core of BRCAS at Academia Sinica.

### snRNA-seq data analysis

#### Data processing and integration

snRNA-seq data from *M. polymorpha* (Tak-1 plants and Mp*hsfb1* mutant) were processed using Seurat (v5.0.0)^56,57^. Raw gene-barcode matrices generated by the 10X Genomics Cell Ranger (v9.0.0) pipeline were imported into R using the Read10X function and converted to Seurat objects. Low-quality nuclei were filtered based on the following criteria: ≥ 250 detected genes, ≥ 500 total unique molecular identifiers (UMIs), and a log_10_ ratio of genes per UMI > 0.8. Following filtering, each dataset was normalized (NormalizeData), highly variable genes were identified (FindVariableFeatures), and expression values were scaled (ScaleData). Principal-component analysis (PCA) was performed, and significant components were selected based on elbow plots. Nuclei were clustered using a shared nearest-neighbor graph (FindNeighbors, FindClusters) and visualized via UMAP (RunUMAP).

Potential multiplets were removed using DoubletFinder (v3.0) with an estimated 7.5% doublet rate^58^. The optimal pK value was selected from the maximum mean-BCmetric score across parameter sweeps, and adjustments were made for homotypic doublets based on cluster composition. Only singlets were retained for downstream analysis. Biological replicates for each genotype (Tak-1 and Mp*hsfb1*) were merged using merge and flattened with JoinLayers to create unified RNA assays. To minimize batch effects, Harmony integration^59^ (RunHarmony) was performed on the first 30 PCs. These integrated embeddings were used for subsequent clustering and visualization.

Finally, the integrated Tak-1 and Mp*hsfb1* objects were aligned. Homologous nuclei were mapped using FindTransferAnchors and TransferData, and cluster correspondences in Mp*hsfb1* were validated using SingleR with the Tak-1 dataset as a reference^60^. Nuclei with a prediction confidence over 0.8 were retained. The final merged object, comprising 18,073 Tak-1 and 14,857 Mp*hsfb1* nuclei, was batch-corrected again using Harmony (grouping by genotype) and visualized by UMAP.

#### Marker gene identification and functional annotation

Cluster-specific marker genes were identified in the integrated Tak-1 and Mp*hsfb1* dataset using FindAllMarkers (Wilcoxon rank-sum test; min.pct = 0.25, logfc.threshold = 0.25, p_adj < 0.01). For each gene, expression specificity was computed as the difference in detection frequency between the target cluster and all other clusters. Marker lists were cross-referenced with a curated *Marchantia* genome (v6.1r2) to annotate transcription factors (TFs). To assign biological functions, cluster markers were converted to *Arabidopsis* orthologs via reciprocal BLAST mapping. GO enrichment analysis was then performed using clusterProfiler (v4.10) with *A. thaliana* TERM2GENE and TERM2NAME mappings^61^. Significantly enriched biological-process terms (FDR < 0.05) were visualized as heatmaps showing GO term count and significance (−log_10_ FDR). Cluster annotation was further refined by matching marker genes to a previously published *Marchantia* single-cell reference^26^. Marker overlap ratios (matched / total annotated markers) were summarized, and the final curated cluster identities were transferred to the Seurat object metadata.

#### Analysis of differentially expressed genes

After integration of Tak-1 and Mp*hsfb1* nuclei, UMAP visualizations and cluster bar plots were generated to assess genotype-specific cell distributions. Cell-type proportions between genotypes were quantified and compared using speckle (v1.1.3) with a logit transformation (propeller)^62^. Differential expression analysis between genotypes was conducted for each annotated cell type using FindMarkers (Wilcoxon test). Genes with an adjusted p < 0.05 and |log_2_ fold-change| > 0.25 were considered significant. DE results were summarized by gene counts and proportions per cell type. To compare transcriptional programs, Jaccard similarity coefficients were calculated between the Tak-1 and Mp*hsfb1* cluster marker sets and visualized as distance heatmaps. Finally, TF expression patterns were summarized by their average log_2_ fold-change across matched cell types.

#### Trajectory inference

Pseudotime trajectories were reconstructed using Monocle3 (v1.3.1)^63-65^ and Slingshot (v2.8.0)^66^ on the integrated Seurat object to capture developmental progression. UMAP coordinates served as the input for trajectory learning; Monocle3 was used to identify the principal graph structure, while Slingshot refined lineage relationships and root assignments. To identify genes dynamically associated with pseudotime, generalized additive models (GAMs) were fitted using tradeSeq (v1.12.0)^67^. The fitGAM function was applied for each lineage from Slingshot, and significant genes were identified via association tests (p < 0.05). Smoothed expression profiles (predictSmooth) were also generated at 100 evenly spaced pseudotime points for each lineage. Genes consistently identified as pseudotime-dependent by both methods were considered trajectory-relevant and used for regulatory network inference.

#### Gene regulatory network (GRN) inference

To ensure regulatory network reconstruction incorporated dynamic genes, the list of pseudotime-associated genes (from Monocle3 + tradeSeq) was merged with expression-based filters before running SCENIC^68^. Specifically, genes expressed in ≥ 1% of nuclei and with total expression > 3% of the mean nucleus count were retained. The union of these filtered genes was the input expression matrix for SCENIC. Regulatory networks (regulons) were inferred independently for Tak-1 and Mp*hsfb1* using SCENIC (v1.3.1) with motif enrichment performed via RcisTarget (v1.29.0)^68^. Gene correlations were computed (runCorrelation), and regulatory links were inferred using GENIE3 (runGenie3(exprMat_filtered, nParts = 20)), partitioning the dataset to optimize memory. This integrated approach ensured that the inferred SCENIC networks reflected genes dynamically modulated along developmental trajectories.

Regulon activity (AUC) matrices in Tak-1 or Mp*hsfb1* SCENIC network were extracted and aligned with cell-type annotations. Average regulon activities and regulon-specificity scores (RSS) per cell type were computed. Genotype-specific and shared regulons were compared using paired Wilcoxon tests, and differential activity was summarized in ΔZ matrices (Tak-1 – Mp*hsfb1*). Network topology metrics—degree, betweenness, closeness, eigenvector, hub, PageRank, and Katz centralities—were computed using igraph (v2.0.3). Global comparisons were performed by Z-score normalization across genotypes, and metric differences were visualized as dot heatmaps, where color represented the ΔZ value and circle size corresponded to magnitude rank. All plots were generated using ggplot2 (v3.5.1) or ComplexHeatmap (v3.21.0)^69^, and all statistical analyses were performed in R (v4.3.1).

### Generation of bulk RNA-seq and bulk RNA-seq analysis

#### Sample collection

For each sample, approximately 200 mg of gemmalings (from 3-, 7-, and 14-day-old plants) or gemmalings with or without prolonged HS treatment (30 °C, 14 days) were promptly transferred to 2 ml Eppendorf tubes, flash-frozen in liquid nitrogen, and stored for subsequent RNA extraction. Total RNA was extracted from the gemmalings using the RNeasy Plant Mini Kit (Qiagen, Hilden, Germany). Library preparation, including polyA enrichment, and sequencing of 150-bp paired-end reads were carried out on the NovaSeq platform at the Genomic Technology Core (IPMB, Academia Sinica, Taiwan), and GENOMICS (Taipei, Taiwan).

#### Data analysis

Raw sequencing reads from *Marchantia* were processed to remove adaptors using fastp (v0.24.0)^70^. The cleaned reads were then mapped to the reference genome (*Marchantia* genome v6.1r2) with STAR (v2.7.9a) using default parameters. Differentially expressed genes (DEGs) were identified using the R package DESeq2^71^, with pairwise comparisons performed against the corresponding genotype at day 3. In the case of prolonged HS, the pairwise comparisons were performed against the corresponding genotype at the control condition. DEGs were defined by a |Log_2_-fold change| ≥ 1.0 and a False Discovery Rate (FDR) < 0.05, using the “lfcShrink ashr” method. Z-scores of expressions were calculated from VST-transformed values. Time-course differential expression analysis was conducted using the splineTimeR (v1.58.0)^72^ package to identify genes with dynamic expression changes across developmental stages. DEGs were identified using the splineDiffExprs function with a smoothing degree of freedom (df) = 2 and a significance threshold of FDR < 0.01. Separate pairwise analyses were performed for Mp*hsfb1* versus Tak-1 and Mp*HSFBox* versus Tak-1 to determine genotype-specific temporal expression patterns. Genes showing significant temporal differences were exported as DEG lists for downstream analyses. Gene Ontology (GO) enrichment analysis was performed using clusterProfiler (v4.10)^61^. Significantly enriched GO terms (FDR < 0.05) were summarized using REVIGO^73^ to reduce redundancy.

### Cell-type-resolved co-expression network analysis using hdWGCNA

#### Data sources and preprocessing

Publicly available single-cell and single-nucleus transcriptomic datasets from Arabidopsis, tomato, rice, and maize (Supporting Dataset S8) were analyzed in R using Seurat for data handling and hdWGCNA (v0.4.0)^74^ for cell-type-resolved co-expression network analysis. Genes were selected for network construction based on expression prevalence to reduce sparsity and technical noise. To improve the robustness of correlation estimates in single-cell data, metacells were generated within each cell type and biological replicate by aggregating transcriptionally similar cells in a low-dimensional space using k = 25 nearest neighbors. Normalized metacell expression profiles were used as input for network inference.

#### Cell-type-specific co-expression network construction

Co-expression networks were constructed independently for each cell type using a signed weighted network framework implemented in hdWGCNA. Soft-thresholding powers were selected based on scale-free topology criteria, requiring a model fit of R² > 0.8. Adjacency matrices were transformed into topological overlap matrices, followed by hierarchical clustering to identify co-expressed gene modules. Module eigengenes were calculated to summarize module-level expression patterns and harmonized across biological replicates where applicable.

#### Quantification of HSFB connectivity

Module connectivity was quantified using eigengene-based connectivity (kME) and node degree for each gene within its assigned module. Genes assigned to the unclassified grey module were excluded from downstream analyses. HSFB-associated modules were defined as modules containing at least one HSFB gene within a given cell type and tissue and were interpreted as putative HSFB co-expressed gene sets. To evaluate the role of HSFBs in each cell type, their connectivity ranks were calculated relative to other transcription factors within the same module and cell type based on kME values. Functional characterization of HSFB-associated modules was further supported by Gene Ontology enrichment analysis using the full hdWGCNA gene set as the background.

### Integration of DAP-seq and scRNA-seq or snRNA-seq datasets

#### Acquisition and preprocessing of DAP-seq datasets

TF binding profiles were obtained from two major DAP-seq resources: GSE60141^33^ and GSE299028^32^. DAP-seq peaks were assigned to target genes if the summit was located within the proximal promoter region, consistently defined as -1000 bp upstream to +500 bp downstream of the annotated transcription start site (TSS). The resulting TF-target lists were harmonized and filtered for unique *Arabidopsis* gene identifiers.

#### Acquisition and preprocessing of scRNA-seq datasets

Single-cell and single-nucleus transcriptomic data were sourced from four *Arabidopsis* atlases: the root atlas^75^ (GSE152766) and the seedling, leaf, and flower bud atlases^32^ (GSE297576). Seurat objects were restricted to the RNA assay and normalized using the default log-normalization method (NormalizeData, LogNormalize). Cell-type annotations were adopted from the original publications.

#### Quantification of TF activity (AUCell)

TF activity was quantified using the AUCell framework^68^, which ranks all genes by normalized expression per cell and computes the area under the recovery curve (AUC) for each DAP-seq gene set. The AUC reflects the enrichment of a TF’s target genes among the most highly expressed genes in that cell. AUC scores were computed from the normalized data layer of each Seurat object using default parameters in AUCell_buildRankings() and AUCell_calcAUC(), producing a cell × TF activity matrix.

#### Cell-type-specific enrichment analysis

TF activity enrichment was identified by comparing AUC scores between cells of a given type and all other cells. Cohen’s *d* (with pooled standard deviation) was calculated using the *effectsize* package to estimate effect size, and one-sided Wilcoxon rank-sum tests assessed directional significance (https://github.com/easystats/effectsize). P-values were adjusted using the Benjamini–Hochberg procedure, and TFs with *d* > 0 and FDR < 0.05 were considered significantly enriched in the corresponding cell type. Results were visualized using dot heatmaps generated

## Data availability

Raw sequencing data have been deposited into the National Center for Biotechnology Information Gene Expression Omnibus and will be made open upon publication.

## Acknowledgment

We thank Ms. Shu-Jen Chou and Ms. Mei-Jen Fang from the Genomics Core at IPMB for preparing the DAP-seq libraries and performing Sanger sequencing, respectively. We are grateful to Dr. Meiyeh Jade Lu (NGS Core Facility, BRC, AS) for generating the snRNA-seq libraries, Dr. Wann-Neng Jane (Cell Biology Core, IPMB) for performing the SEM experiments, and Dr. Yu-Chih Lin (Metabolomics Core, ABRC, AS) for measuring cytokinin concentrations. We also acknowledge the Flow Cytometry Cores at IPMB and IBMS, AS, for assistance with nuclei sorting, as well as the Bioinformatics Core at IPMB for high-performance computing support. Furthermore, we thank Drs. Chih-Hang Wu and Shu-Hsing Wu (IPMB, AS) for providing plasmids for the PPI and LUC reporter assay, and Dr. Yucheng Long (National University of Singapore) for helpful discussions on quantifying solidity using ImageJ.

## Author’s contribution

TYW conceptualized and oversaw all the experiments and analyses. HCT and KHL designed and performed all the experiments with the assistance of HJY and TYW. YRL and MJL performed PPI experiments. LT performed snRNA-seq and downstream analysis. MRY and TYW performed formal bioinformatic analyses with assistance from CYC. TYW wrote the manuscript with assistance from all authors. HCT, KHL, CYC, and TYW edited the manuscript.

## Supplemental information

**Supporting Figure S1.**
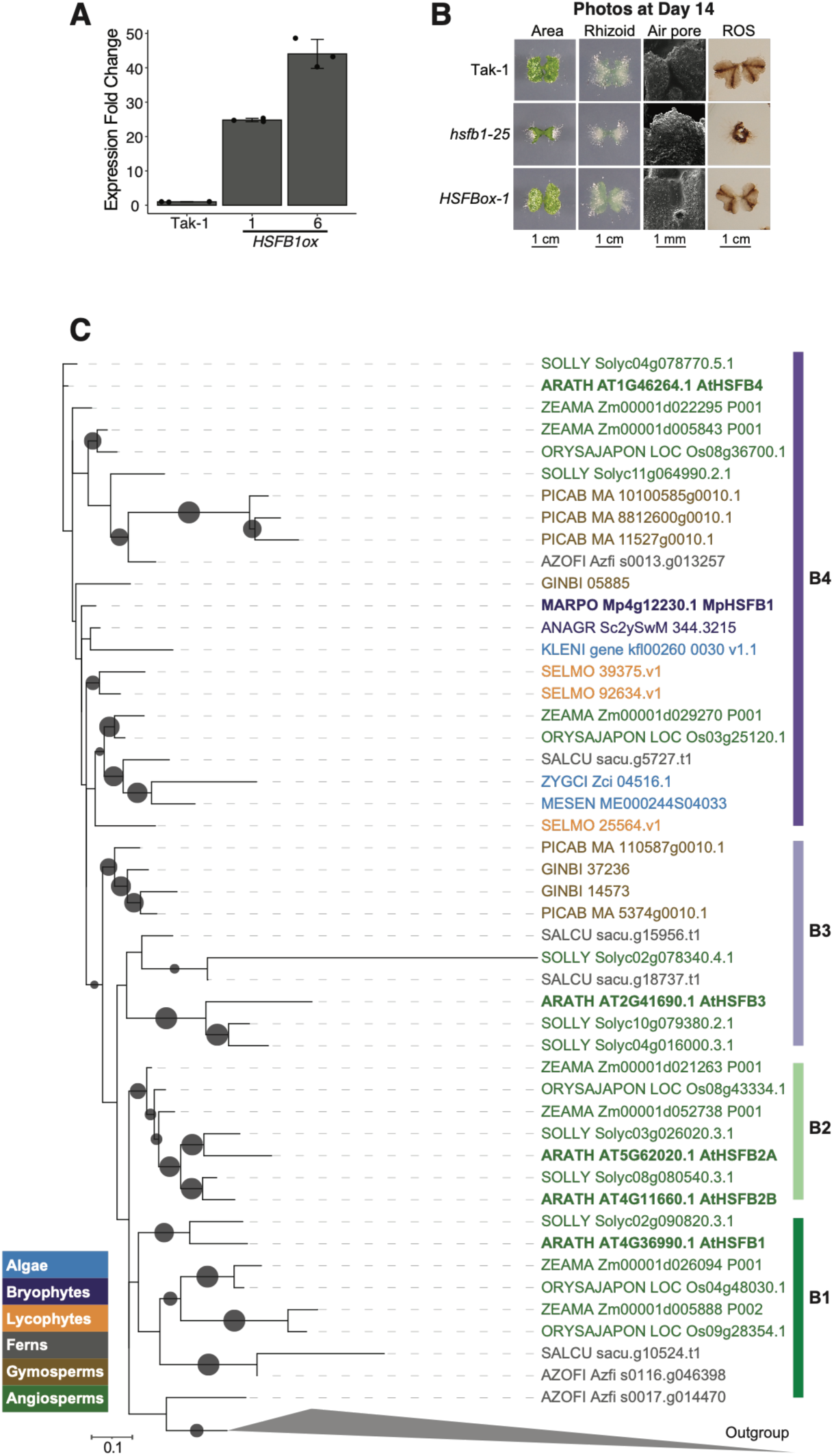
Characterization of Mp*hsfb1* knockout and Mp*HSFB1ox* overexpression lines. **A.** Bar graph showing MpHSFB1 expression levels in Tak-1 plants, as well as Mp*HSFB1ox-1* and Mp*HSFB1ox-6* lines. Fold change was calculated relative to endogenous MpHSFB1 expression in Tak-1. *N* = 3 biological replicates. **B.** Representative photos of the second independent lines of Mp*hsfb1* and Mp*HSFB1ox* plants. Note that the photo for Tak-1 plants is from the same set as in Figure 1, as the experiments were performed simultaneously. **C.** Phylogenetic tree of plant HSFB sequences. The maximum-likelihood tree was constructed using the DNA-binding domain (DBD) sequences of selected taxa. The HSFB1–B4 groups are color-coded, and all sequences used for tree construction are listed in Supporting Dataset 1.

**Supporting Figure S2.**
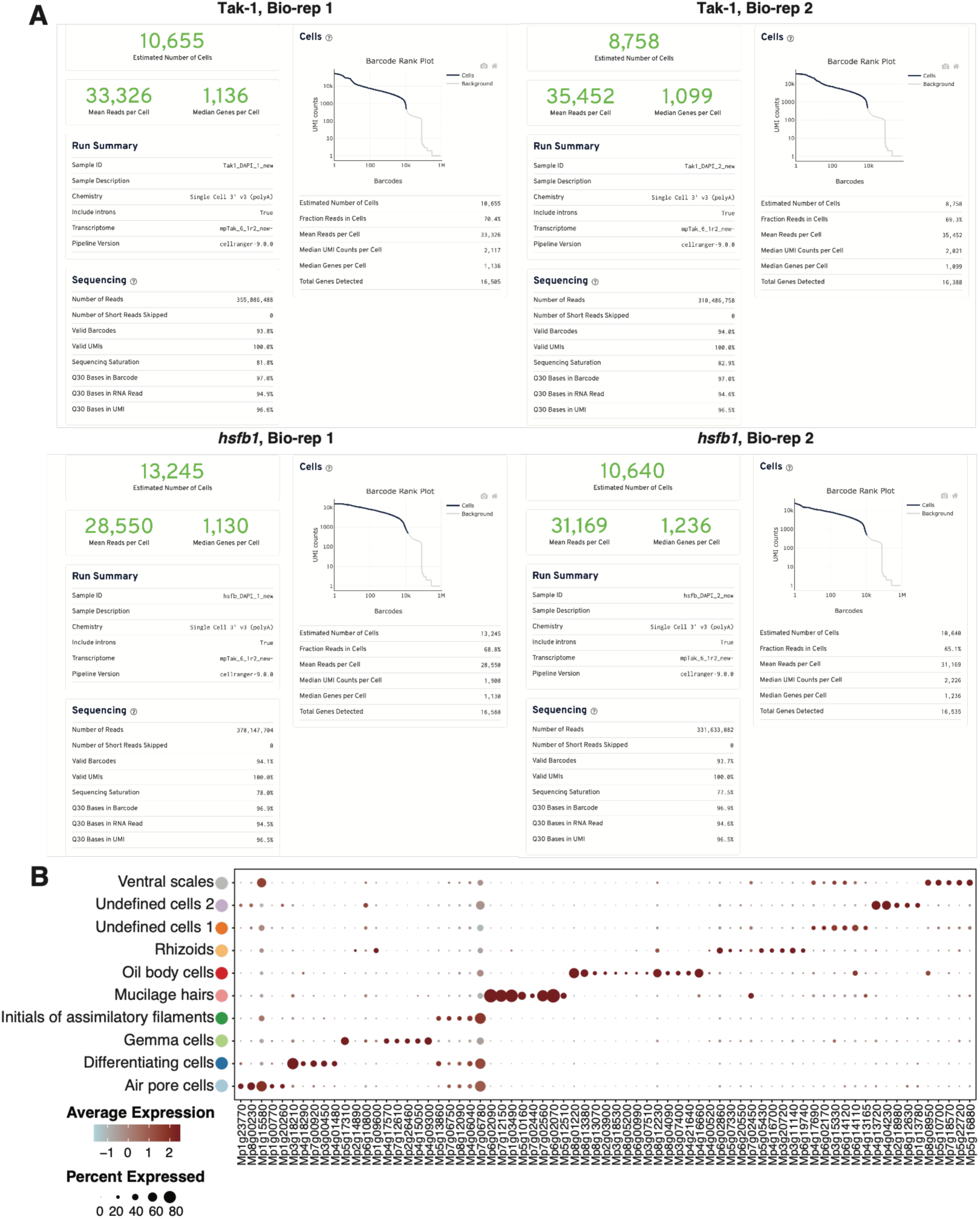
Quality control of snRNA-seq data and identification of cell-type-specific marker genes. **A.** Output files from the Cell Ranger software for each snRNA-seq sample. **B.** Dot plots showing the marker genes for each known cell type. Information regarding marker genes and cell types was taken from a previous study^26^. Dot size indicates the percentage of cells expressing the gene (% expressed). Dot colors represent the proportional amount of expression of each gene in each cell type, with deeper colors indicating higher expression levels.

**Supporting Figure S3.**
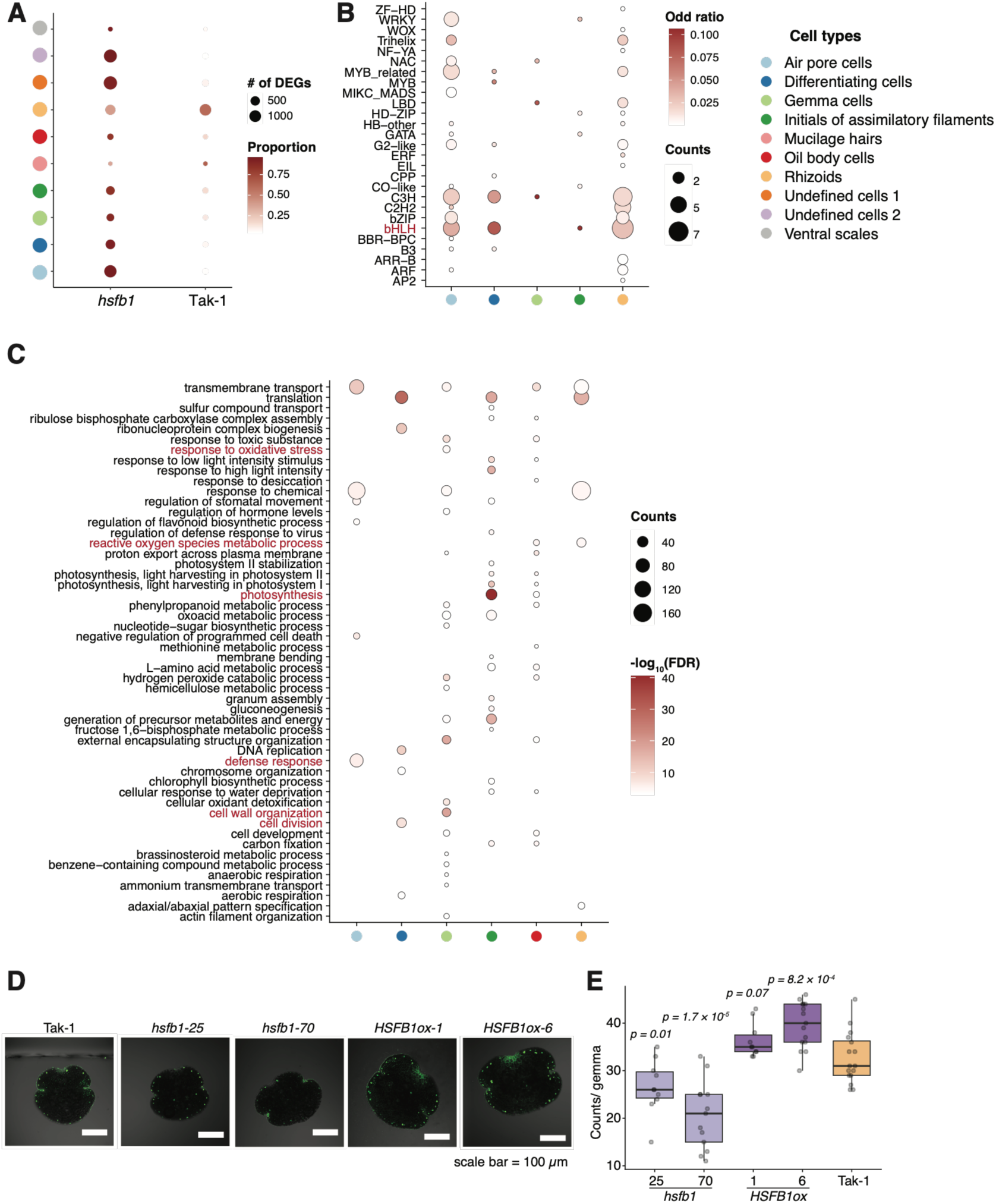
Characterization of cell-type-specific DEGs in Tak-1 plants and Mp*hsfb1* mutants. **A.** Dot plot summarizing the proportion of DEGs found in each cell type in Tak-1 plants and Mp*hsfb1* mutants. **B.** Dot plot showing the counts of differentially expressed transcription factors (DETFs) in each cell type. The color represents the odds ratio. **C.** Dot plot showing the overrepresented GO terms in each cell type. The size of the circle represents the DEG count. **D.** Representative images of oil body staining for each genotype. **E.** Boxplot showing the number of oil bodies per gemma for each genotype. Boxplots display all individual data points (dots), median values, first and third quartiles, and whiskers indicating minimum and maximum values. *P*-values were determined by a pairwise Student’s *t*-test. *n* = 10.

**Supporting Figure S4.**
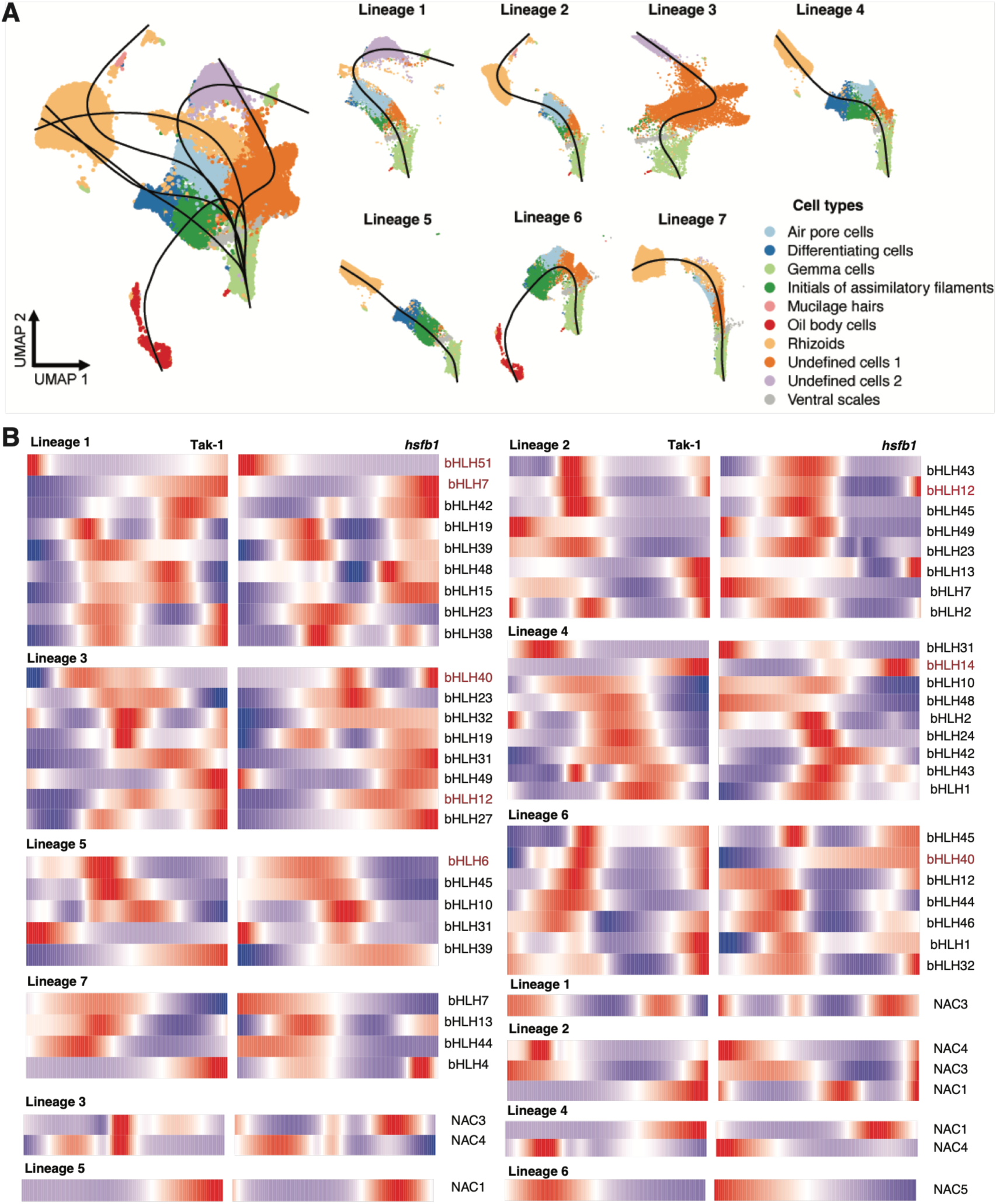
Pseudotime trajectory and cell lineage analysis in snRNA-seq. **A.** UMAP showing the lineage analysis from Slingshot. In total, seven possible lineages were identified. **B.** Heatmap showing the expression patterns of differentially expressed bHLH and NAC TFs in each lineage within the Tak-1 and Mp*hsfb1* datasets. The color bar indicates the relative expression level.

**Supporting Figure S5.**
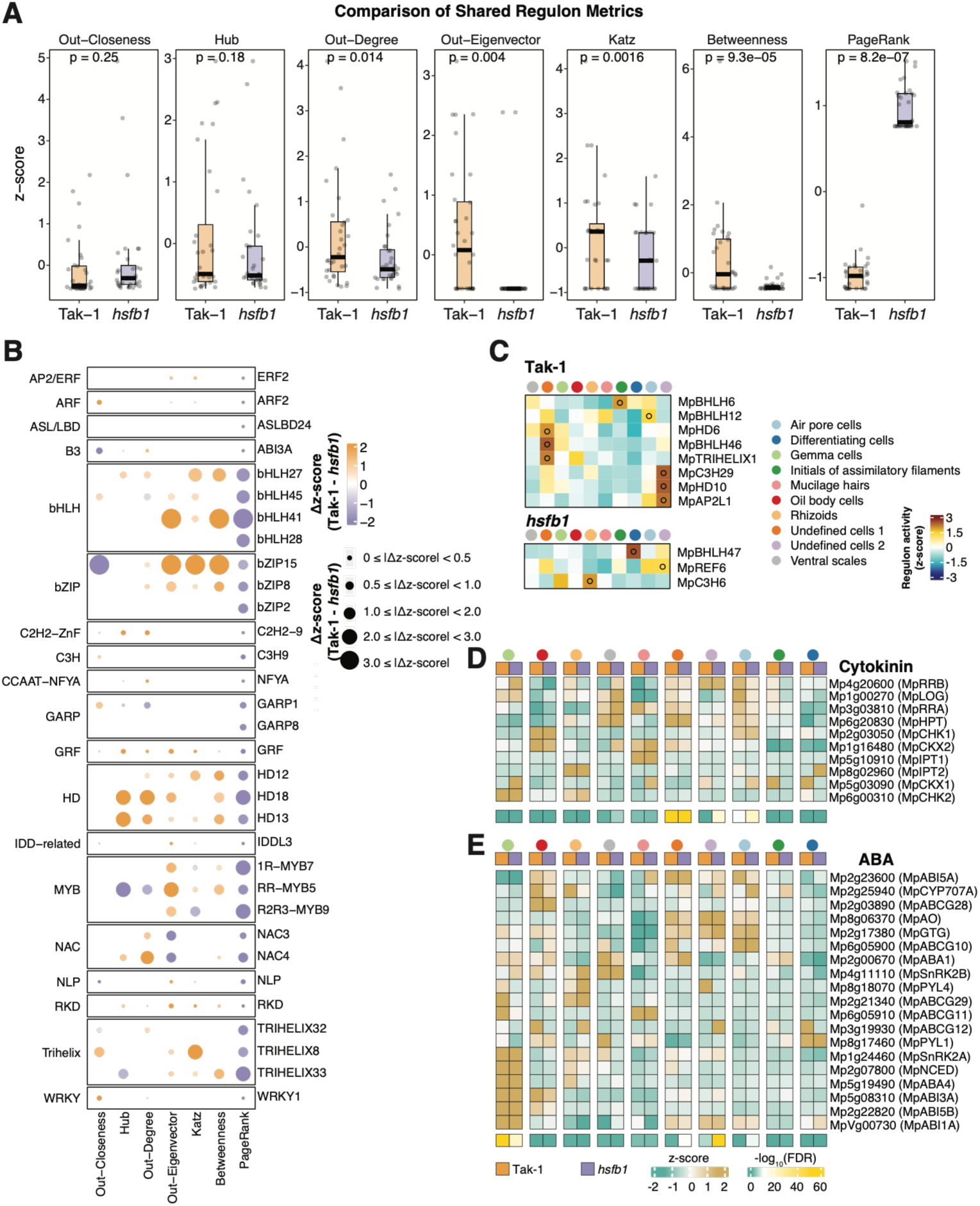
snRNA-seq network analysis and identification of hormone-related gene expression. **A.** Boxplots showing seven metrics of shared regulons identified from Tak-1 and Mp*hsfb1* mutants. Dots represent individual plant data; p-values were determined by the Mann-Whitney U test with Bonferroni correction. Boxplots display all data points (dots), median values, first and third quartiles, and whiskers indicating minimum and maximum values. **B.** Dot plots showing the scaled regulon specificity score (RSS) after z-score transformation in Tak-1 and Mp*hsfb1* for each network metric. Dot size indicates the value of Δz-score, and dot color denotes the Δz-score between Tak-1 and Mp*hsfb1*. **C.** Heatmap showing the genotype-specific scaled regulon specificity score (RSS) after z-score transformation in Tak-1 and Mp*hsfb1*. The color bar represents the activity of regulons. **D & E.** Heatmaps showing relative expression of genes related to cytokinin (**D**) and abscisic acid (ABA) (**E**) biosynthesis and signaling in each annotated cell type and genotype. Gene expression levels are represented by aggregated unique molecular identifier (UMI) counts across all cells belonging to the same cell type within one genotype. The adjusted p-value was determined by the hypergeometric distribution.

**Supporting Figure S6.**
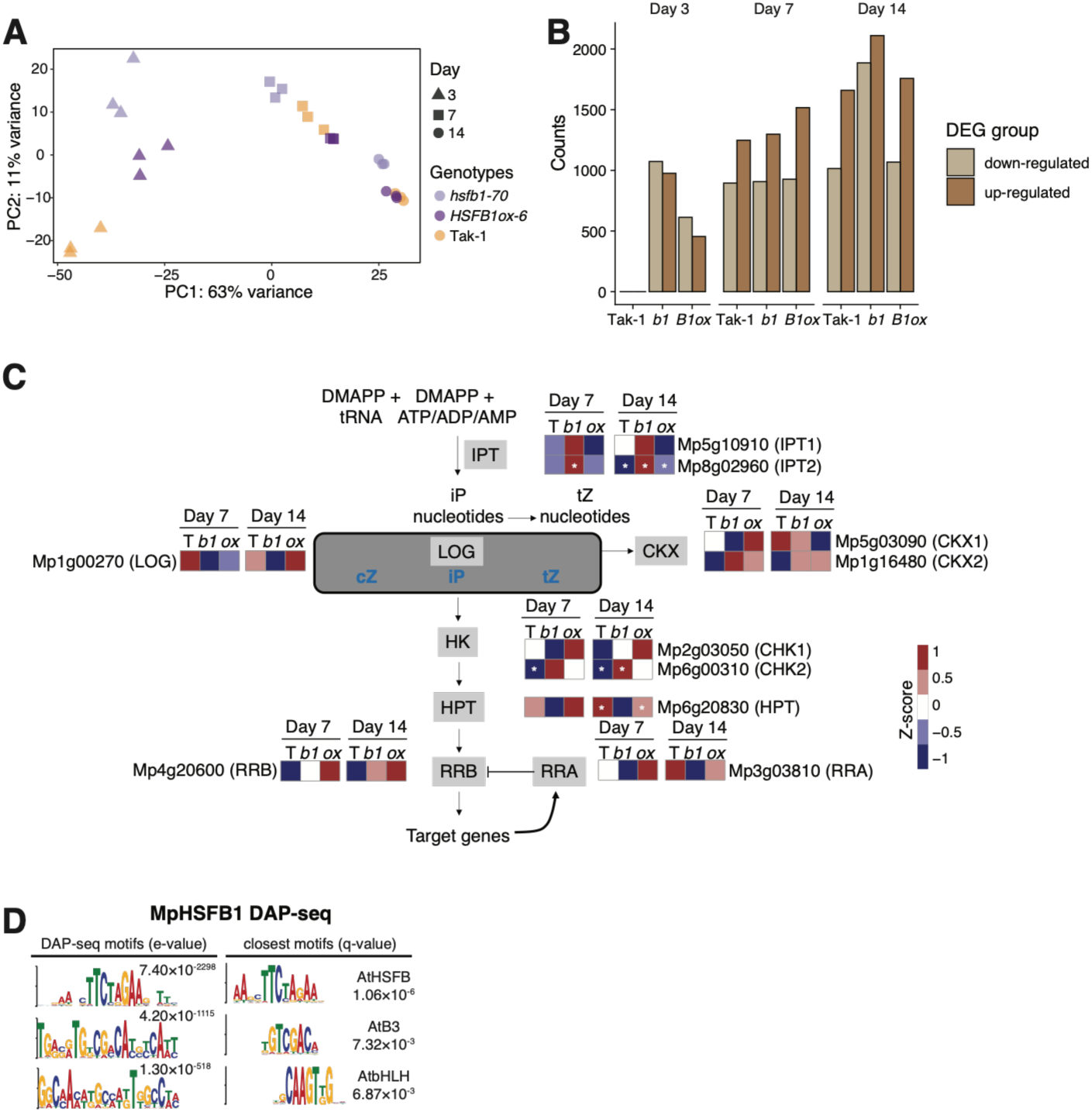
Time-course bulk RNA-seq analysis of Mp*hsfb1*, Mp*HSFB1ox*, and Tak-1 plants. **A.** Principal Component Analysis (PCA) comparing RNA-seq biological replicates from each genotype at days 3, 7, and 14. **B.** Bar graph displaying the number of differentially expressed genes (DEGs) in each genotype at days 3, 7, and 14. Note that data from Tak-1 plants at day 3 were used as the baseline for pairwise comparisons between mutants and wild-type plants; therefore, no DEGs are shown for Tak-1 at day 3. For comparisons at day 7 and day 14, the respective genotype’s data at day 3 were used as the baseline. **C.** Heatmap showing the expression profiles of cytokinin biosynthesis genes in Tak-1, Mp*HSFB1-ox*, and Mp*hsfb1* plants at days 7 and 14 relative to day 3. Asterisks indicate significant DEGs (|log2FC| ≥ 1, FDR < 0.01). **D.** Top three binding motifs determined by DAP-seq and their closest known TFBSs.

**Supporting Figure S7.**
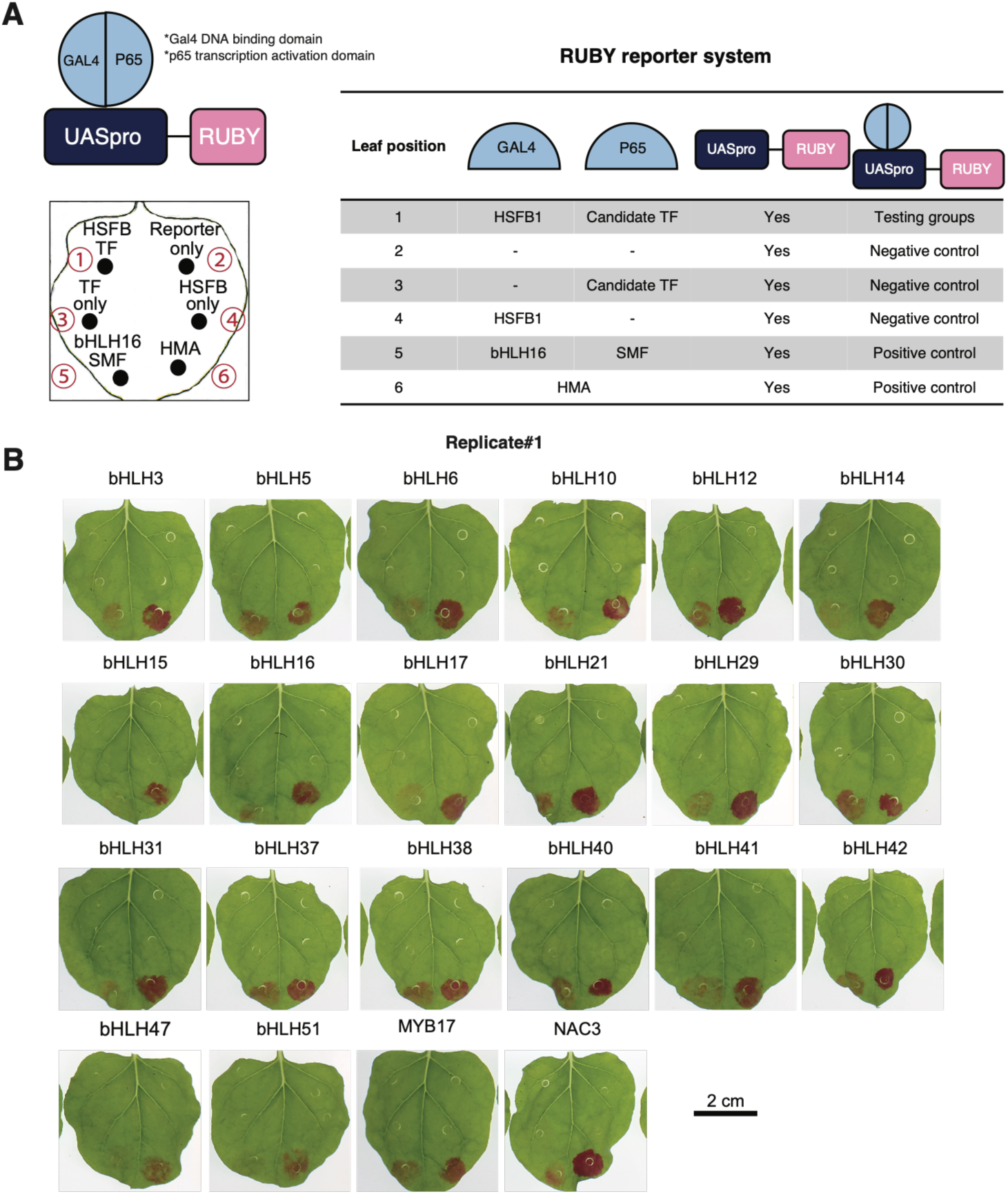
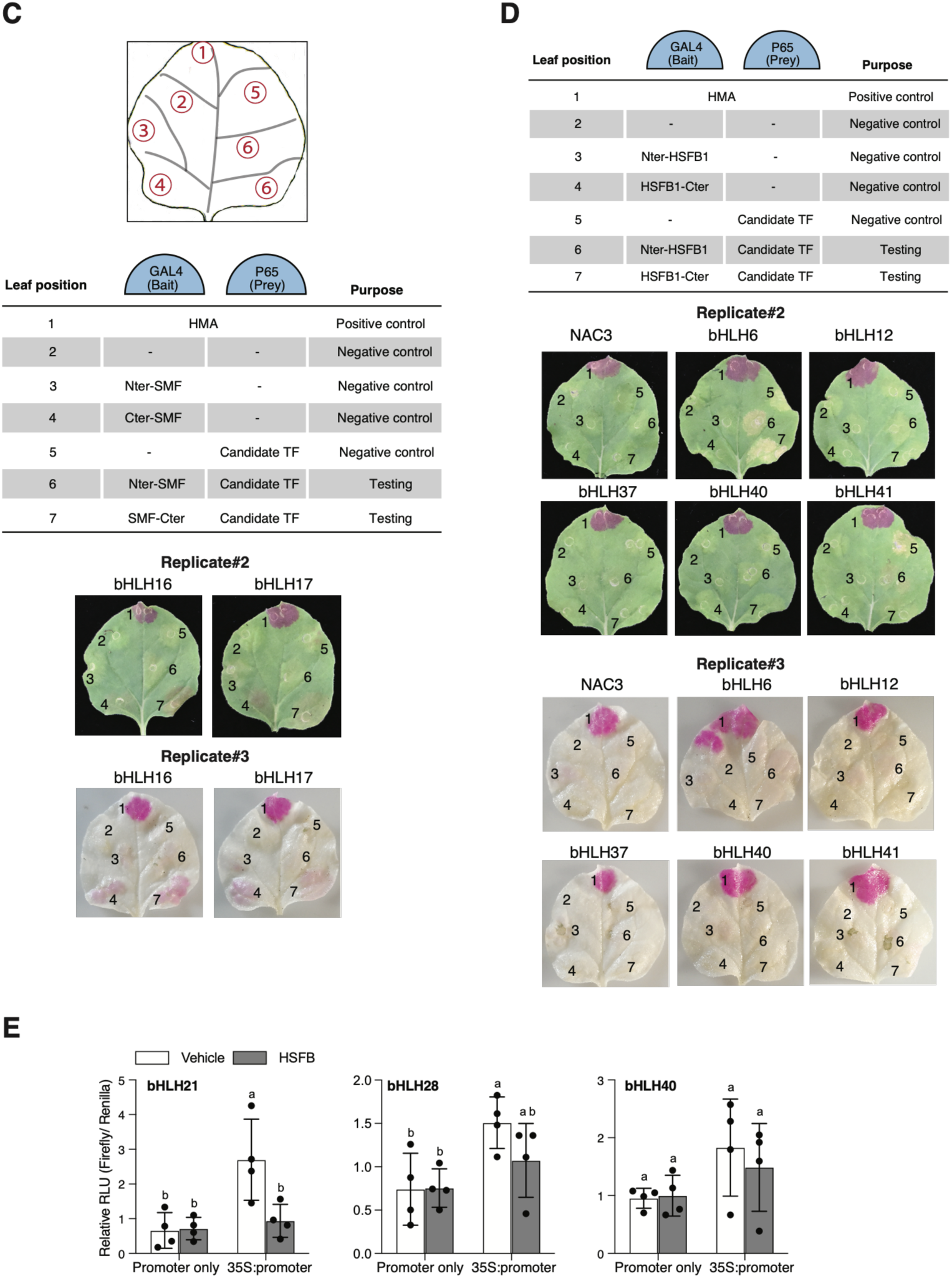
Validation of MpHSFB1–MpbHLH protein-protein interactions in tobacco leaves. **A.** Schematic illustrating construct designs for protein-protein interaction using the split GAL4-RUBY assay and the arrangement of infiltration sites on tobacco leaves. The known interaction between bHLH16 and SMF served as the positive control^76^. **B.** Representative photos showing betalain production in tobacco leaves for each tested TF. Experiments were repeated three times with three leaves per replicate, yielding consistent results. **C. & D**. Independent biological replicates (second and third) of the experiments shown in A and B. Note that in the third biological replicate, leaves were destained with 95% ethanol to remove chlorophyll for better visualization. **E.** Dual-luciferase reporter activity (firefly/*Renilla*) measured in *N. benthamiana* co-transfected with each *MpBHLHpro:firefly* reporter and either vehicle (white bars) or the MpHSFB1 effector (grey bars). For each reporter, activity is shown under the reporter alone (Promoter only) and with the effector driven by the CaMV 35S promoter (35S:promoter). Bars represent mean ± SE (n = 4 biological replicates). Different letters denote statistically significant differences (p < 0.05, two-way ANOVA with Tukey’s post-hoc test).

**Supporting Figure S8.**
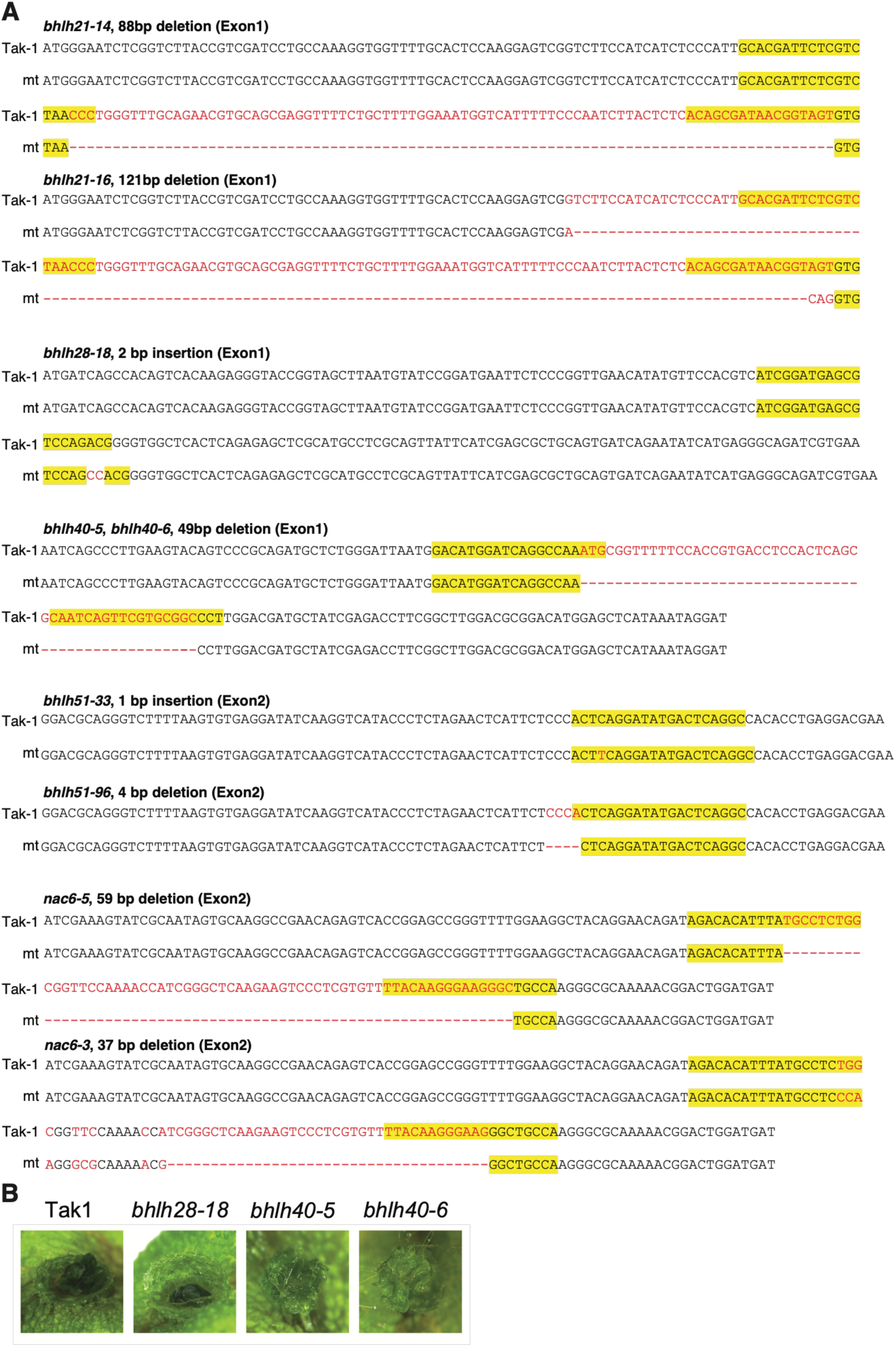
Characterization of Mp*bhlh* knockout mutants. **A.** Indel sequences of Mp*bhlh21-14*, Mp*bhlh21-16*, Mp*bhlh28-18*, Mp*bhlh40-5*, Mp*bhlh40-6*, Mp*bhlh51-33*, Mp*bhlh51-96*, Mp*nac6-3*, and Mp*nac6-5* plants. Red text indicates deletions, and yellow boxes highlight the gRNA target sites. **B.** Representative photos illustrating gemma cup morphology in Mp*bhlh28-18*, Mp*bhlh40-5*, and Mp*bhlh40-6* plants compared to Tak-1 plants under control conditions. Photos were taken after 24 days of growth on ½ × B5 plates.

**Supporting Figure S9.**
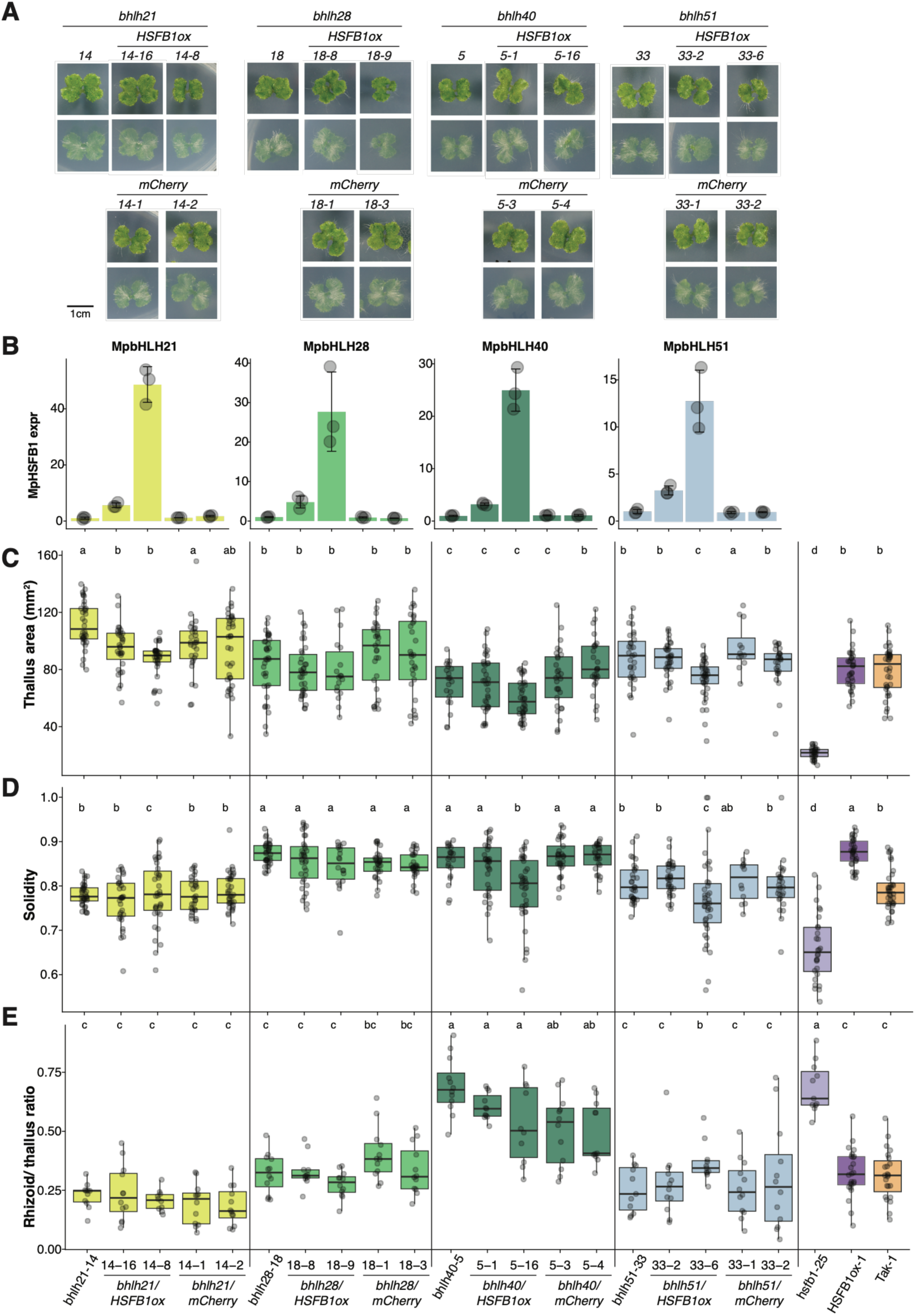
Characterization of Mp*bhlh*/Mp*HSFB1ox* lines. **A.** Representative images of Mp*bhlh*/Mp*HSFB1ox* lines and their corresponding vector controls after 14 days of growth on ½ × B5 plates. **B.** Relative *MpHSFB1* expression levels measured by RT-qPCR across all transgenic lines shown in (**A**). **C.–E.** Boxplots showing thallus area (**C**), solidity (**D**), and rhizoid-to-thallus ratio (**E**) for each genotype. Lowercase letters indicate statistically significant differences determined by one-way ANOVA with Tukey’s post hoc HSD test. For all panels, boxplots display individual data points (dots), median values, first and third quartiles, and whiskers indicating minimum and maximum values.

**Supporting Figure S10.**
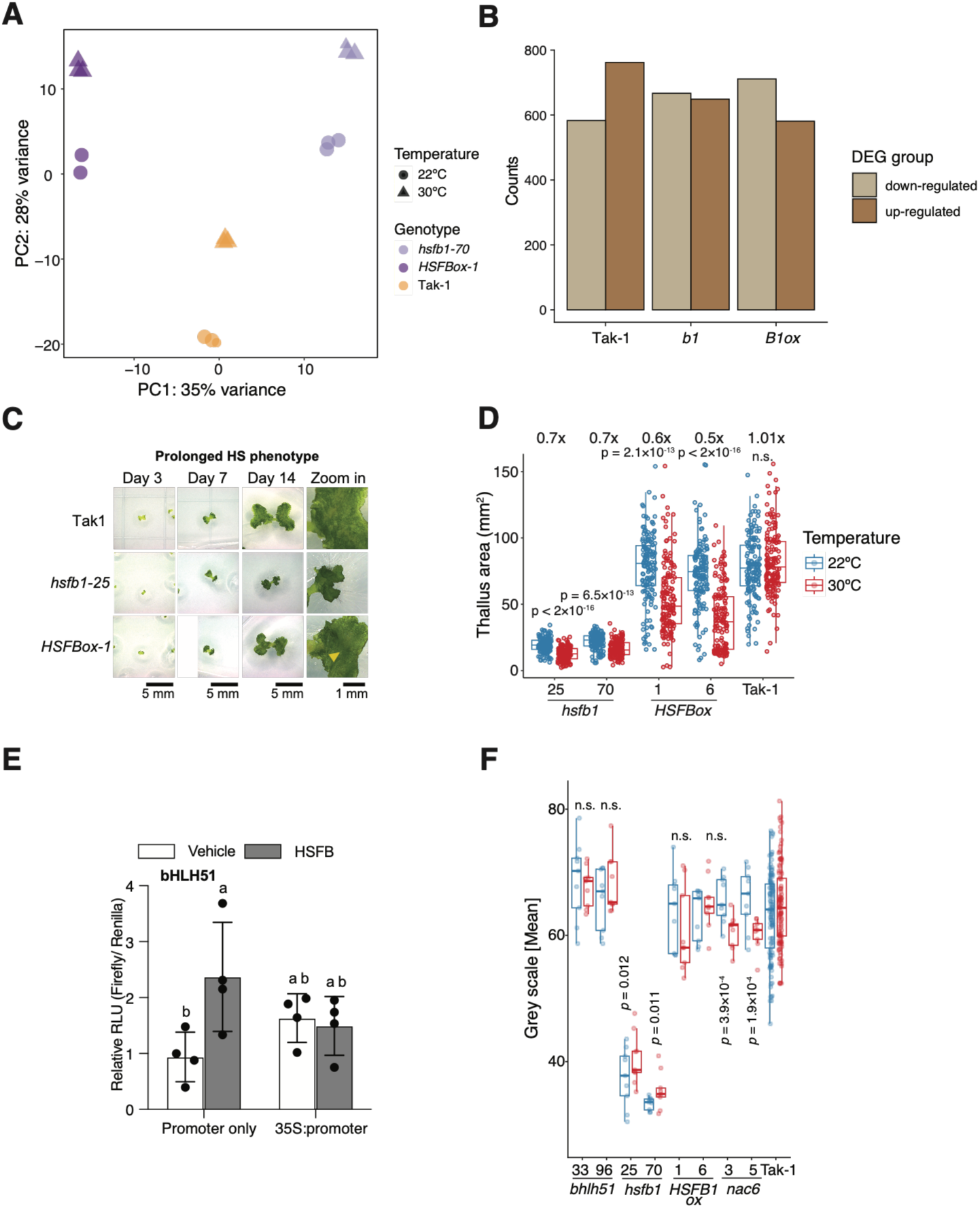
Analysis of bulk RNA-seq datasets in Mp*hsfb1*, MpH*SFB1ox*, and Tak-1 plants under prolonged HS. **A.** Principal Component Analysis (PCA) comparing RNA-seq biological replicates from each genotype under each condition. **B.** Bar graph displaying the number of DEGs in each genotype under prolonged HS treatment. **C.** Representative photos of the second independent lines of Mp*hsfb1* and Mp*HSFB1ox* plants under prolonged HS treatment. Note that the photo for Tak-1 plants is from the same set as in Figure 5A (HS condition), as the experiments were performed simultaneously. **D.** Boxplots showing thallus area (in pixels) for each genotype and condition. P-values were determined by the Mann-Whitney U test with Bonferroni correction (*n* = 60 plants per line). Fold change was calculated as the ratio of the mean thallus area between control and HS treatments for each genotype. **E.** Dual-luciferase reporter activity (firefly/Renilla) measured in *N. benthamiana* co-transfected with each *MpBHLHpro:firefly* reporter and either vehicle (white bars) or the MpHSFB1 effector (grey bars). For each reporter, activity is shown under the reporter alone (Promoter only) and with the effector driven by the CaMV 35S promoter (35S:promoter). Bars represent mean± SE (n = 4 biological replicates). Different letters denote statistically significant differences (p < 0.05, two-way ANOVA with Tukey’s post-hoc test). **F.** Boxplots showing DAB staining intensities in each genotype and condition. The y-axis denotes grayscale pixel values (0 [black]–255 [white]). P-values were determined by the Mann-Whitney U test with Bonferroni correction (*n* = 60-100 plants per line). For all panels, boxplots display individual data points (dots), median values, first and third quartiles, and whiskers indicating minimum and maximum values.

**Supporting Figure S11.**
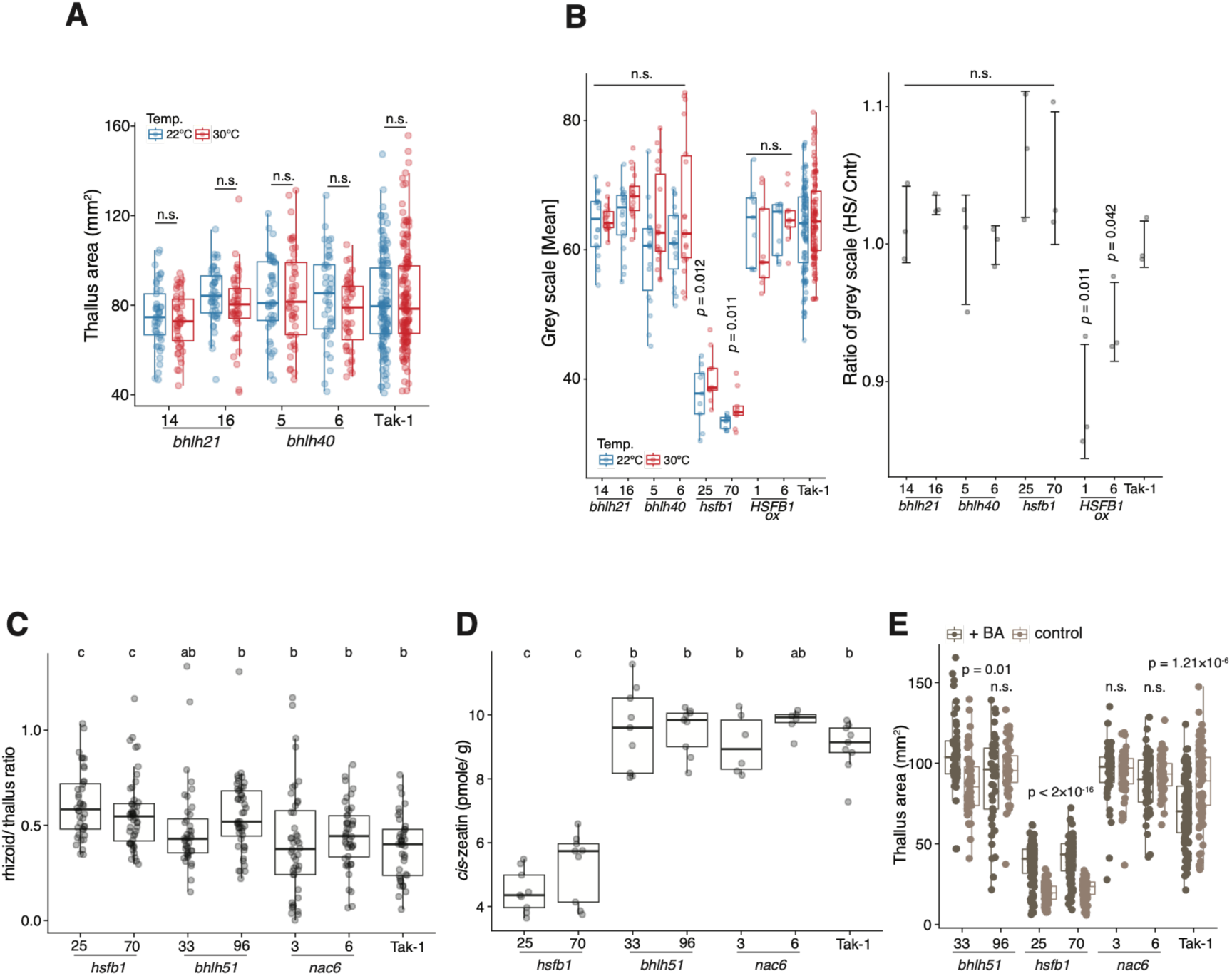
Regulatory specificity of MpHSFB1-mediated MpbHLH regulation. **A.** Boxplots showing thallus area for each genotype and condition. Dots represent individual plant data; *p*-values were determined by the Mann-Whitney *U* test with Bonferroni correction (*n* = 60 plants per line). **B.** (Left) Boxplots showing DAB staining intensities for each genotype and condition. The y-axis denotes grayscale pixel values from 0 (black) to 255 (white). P-values were determined by the Mann-Whitney *U* test with Bonferroni correction (*n* = 12–60 plants per line). (Right) Dot plot showing the grayscale ratio under prolonged HS relative to control conditions for each genotype. Error bars indicate the standard error (S.E.), where a lower ratio indicates higher ROS levels. **C.** Boxplot showing the rhizoid-to-thallus ratio. P-values were determined by one-way ANOVA followed by Tukey’s HSD *post hoc* test (*n* = 10–12 plants per line). **D.** Boxplot showing *cis*-zeatin levels. P-values were determined by one-way ANOVA followed by Tukey’s HSD *post hoc* test (*n* = 6–8 independent measurements). **E.** Boxplots showing thallus area for each genotype and condition. P-values were determined by the Mann-Whitney *U* test with Bonferroni correction (*n* = 30–60 plants per line). For all panels, boxplots display individual data points (dots), median values, first and third quartiles, and whiskers indicating minimum and maximum values.

**Supporting Figure S12.**
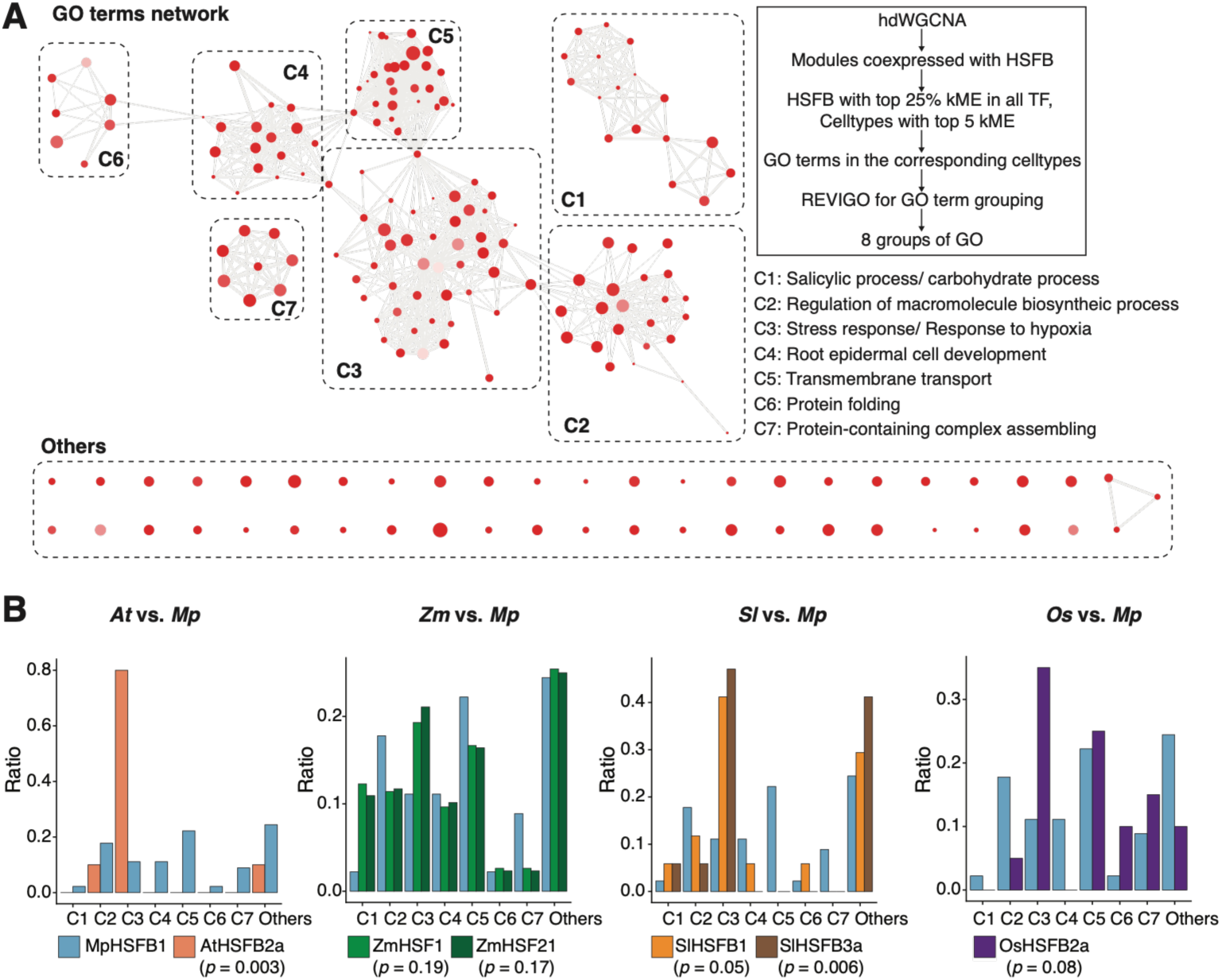
HSFB co-expression networks identified by hdWGCNA across various land plants. (A) Network showing 8 groups of GO term categories with their corresponding functions. The color of the node represents the significance of the GO biological process (BP) term (-log10(FDR)), and the size of the node represents the total number of annotated genes. (B) Bar graphs showing the proportion of GO terms in each GO category for selected HSFBs. P-values were determined by Fisher’s exact test (comparisons made against MpHSFB1).

**Supporting Figure S13.**
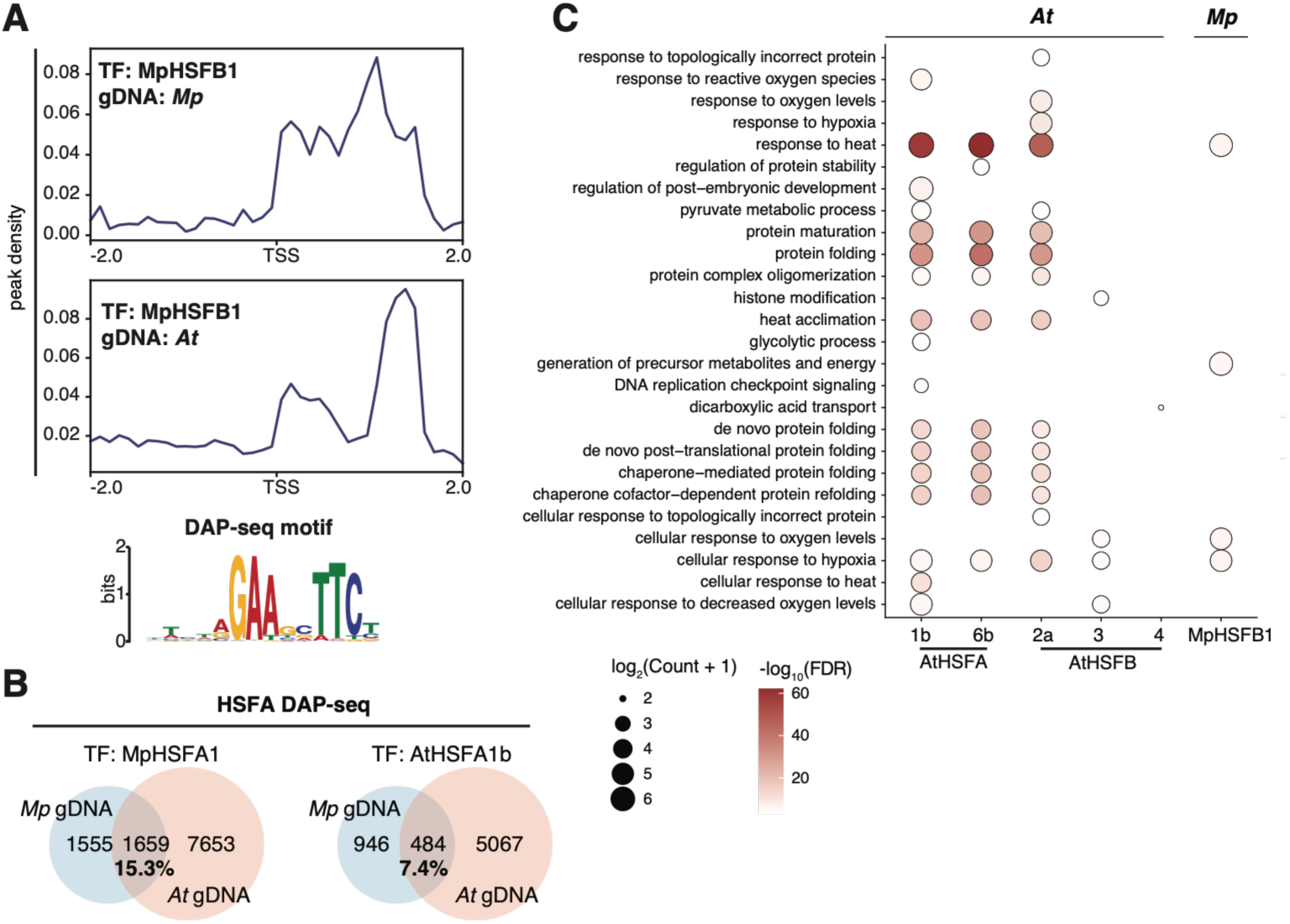
Comparative swapDAP-seq analysis in *Marchantia* and *Arabidopsis*. **A.** Metaplots showing the distribution of MpHSFB1 on *Marchantia* and *Arabidopsis* genomic DNA in the region (± 2 kb) surrounding transcription start sites (TSSs). The sequence logo of the binding motif identified from *Arabidopsis* genomic DNA is shown in the bottom panel. **B.** Venn diagrams showing overlapping orthogroups identified by swap-DAP-seq using MpHSFA1 or AtHSFA1b bound to the *Marchantia* and *Arabidopsis* genomes. **C.** Dot plot showing overrepresented GO terms in each cell type. Circle size represents the count of target genes for each HSF.

**Supporting Figure S14.**
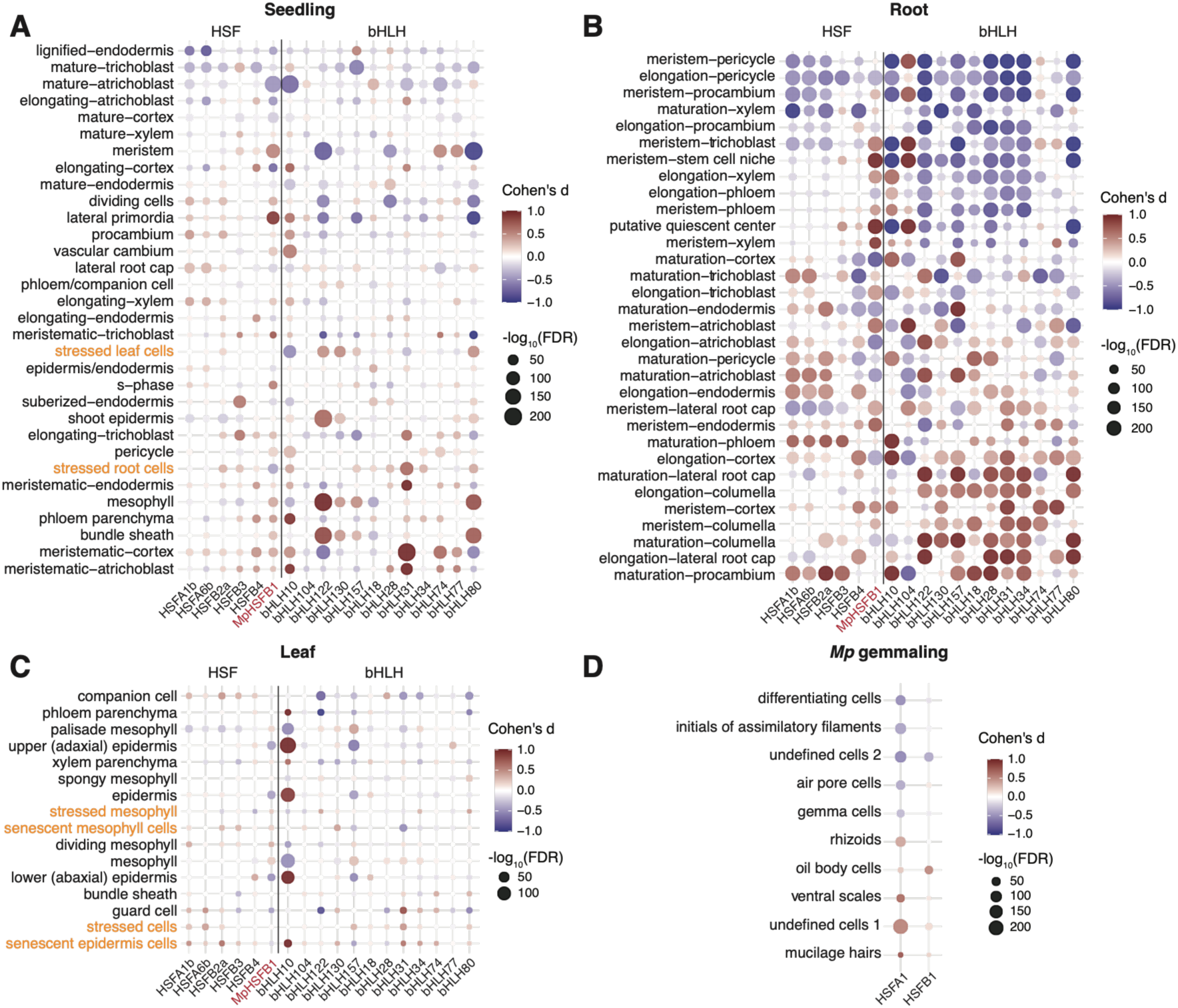
Multi-tissue regulatory map of HSFB- and bHLH-mediated cell-type-specific gene expression programs. **A–C.** Dot plots showing the enrichment or depletion of target gene expression for each HSFA and HSFB (horizontal axis) across labeled cell types in the Arabidopsis root (**A**), seedling (**B**), and flower bud (**C**) snRNA-seq atlases (vertical axis). Dot size represents the FDR-corrected −log_10_(p-value) from a Wilcoxon rank-sum test of summarized target expression scores for each cell-type label versus all other cells. Dot color represents the effect size (Cohen’s d) of expression enrichment (red) or depletion (blue). Enrichments with FDR-corrected p < 0.01 are shown. **D.** Same as A–C but for the Marchantia snRNA-seq atlas.

**Supporting Dataset S1. Raw phenotypic data under control, cytokinin, and prolonged HS conditions, and endogenous cytokinin measurements.**

**Supporting Dataset S2. Quality control of snRNA-seq data, cell type annotation, and identification of differentially expressed genes.**

**Supporting Dataset S3. Results of cell lineage analysis using Slingshot.**

**Supporting Dataset S4. snGRN analysis in Tak-1 and Mp*hsfb1* plants.**

**Supporting Dataset S5. GO enrichment analysis results for each regulon.**

**Supporting Dataset S6. List of DEGs from 3-, 7-, and 14-day-old Tak-1, Mp*hsfb1*, and Mp*HSFB1ox* plants, and list of MpHSFB1-targeted genes from DAP-seq.**

**Supporting Dataset S7. Lists of DEGs from 7- and 14-day-old Mp*bhlh21* and Mp*bhlh40* mutants, alongside re-analyses of publicly available Mp*bHLH* mutant RNA-seq datasets.**

**Supporting Dataset S8. List of DEGs from Tak-1, Mp*hsfb1*, Mp*HSFB1ox,* Mp*bhlh51 and* Mp*nac6* plants under prolonged HS.**

**Supporting Dataset S9. List of co-expression modules and corresponding GO terms identified by hdWGCNA in *Arabidopsis*, rice, tomato, maize, and *Marchantia* scRNA-seq or snRNA-seq datasets.**

**Supporting Dataset S10. List of genes identified from swapDAP-seq and HSFB regulons identified from different scRNA-seq or snRNA-seq datasets.**

**Supporting Dataset S11. List of primers and plasmids used in this study.**

